# Systems immune profiling of variant-specific vaccination against SARS-CoV-2

**DOI:** 10.1101/2021.12.02.471028

**Authors:** Lei Peng, Jonathan J. Park, Zhenhao Fang, Xiaoyu Zhou, Matthew B. Dong, Qiancheng Xiong, Chenxiang Lin, Sidi Chen

## Abstract

Lipid-nanoparticle(LNP)-mRNA vaccines offer protection against COVID-19. However, multiple variant lineages caused widespread breakthrough infections. There is no report on variant-specific vaccines to date. Here, we generated LNP-mRNAs specifically encoding wildtype, B.1.351 and B.1.617 SARS-CoV-2 spikes, and systematically studied their immune responses in animal models. All three LNP-mRNAs induced potent antibody responses in mice. However, WT-LNP-mRNA vaccination showed reduced neutralization against B.1.351 and B.1.617; and B.1.617-specific vaccination showed differential neutralization. All three vaccine candidates elicited antigen-specific CD8 and CD4 T cell responses. Single cell transcriptomics of B.1.351-LNP-mRNA and B.1.617-LNP-mRNA vaccinated animals revealed a systematic landscape of immune cell populations and global gene expression. Variant-specific vaccination induced a systemic increase in reactive CD8 T cell population, with a strong signature of transcriptional and translational machineries in lymphocytes. BCR-seq and TCR-seq unveiled repertoire diversity and clonal expansions in vaccinated animals. These data provide direct systems immune profiling of variant-specific LNP-mRNA vaccination *in vivo*.

## Introduction

Severe acute respiratory syndrome coronavirus (SARS-CoV-2), the pathogen of coronavirus disease 2019 (COVID-19), has caused the ongoing global pandemic (Dong et al., 2020). Although lipid nanoparticle (LNP) - mRNA based vaccines such as BNT162b2 (Pfizer-BioNTech) and mRNA-1273 (Moderna) have demonstrated high efficacy against COVID-19, breakthrough infections have been widely reported in fully vaccinated individuals(Hacisuleyman et al., 2021; Kustin et al., 2021) (Bergwerk et al., 2021; Callaway, 2021a; Mizrahi et al., 2021; Stephen J. Thomas, 2021; Tyagi et al., 2021). Moreover, the virus continues to mutate and multiple dangerous variant lineages have evolved, such as B.1.1.7, B.1.351, and, more recently B.1.617 (Liu et al., 2021; Taylor et al., 2021b). The B.1.1.7 lineage (Alpha variant, or “UK variant”) has an increased rate of transmission and higher mortality (Davies et al., 2021). The B.1.351 lineage (Beta variant, or “South Africa variant”) has an increased rate of transmission, resistance to antibody therapeutics, and reduced vaccine efficacy (Tegally et al., 2021; Wang et al., 2021b; Zhou et al., 2021a). The lineage B.1.617 (“Indian variant” lineage, including B.1.617.1 “Kappa variant”, B.1.617.2 “Delta variant” and B.1.617.3) has recently emerged, spread rapidly, and become dominant in multiple regions in the world (Rambaut et al., 2020, 2021). The on-going surge of infections in the US is predominantly caused by the Delta variant, originating from the B.1.617 lineage that has >1,000 fold higher viral load in infected individuals (Baisheng Li, 2021; Reardon, 2021). The B.1.617 lineage has an increased rate of transmission, showing reduced serum antibody reactivity in vaccinated individuals, and exhibits resistance to antibody therapeutics (Planas et al., 2021a; Singh et al., 2021; Tada et al., 2021; Wall et al., 2021; Yadav et al., 2021). More recently, the Omicron variant (B.1.1.529), representing the most heavily mutated variant of SARS-CoV-2 identified to date, emerged and rapidly spread across the global (Callaway, 2021b) (Pulliam et al. 2021) (https://www.gisaid.org). All these variants often spread faster than the original “wildtype” (WT) virus (also noted as Wuhan-Hu-1 or WA-1). Several of them are known to cause more severe disease, are more likely to escape certain host immune response, cause disproportionally higher numbers of breakthrough infections despite the status of full vaccination (Abu-Raddad et al., 2021; Lopez Bernal et al., 2021a; Tegally et al., 2021; Wang et al., 2021a; Zhou et al., 2021b), and have been designated by WHO and CDC as “variants of concern” (VoCs) (Taylor et al., 2021a). Regarding their effects on vaccine efficacy, B.1.351, for example, has been known to reduce the efficacy of the Pfizer-BioNTech vaccine from >90% to near 70%(Abu-Raddad et al., 2021). The Delta variant has also resulted in significant reduction of vaccine efficacy especially for individuals who received only a single dose (Lopez Bernal et al., 2021a), and has caused wide-spread breakthrough infections despite the status of full vaccination(Brown et al., 2021).

It has been widely hypothesized that the next-generation of COVID-19 vaccines can be designed to directly target these variants (“variant-specific vaccines”). However, to date, there has been no literature report on any approved or clinical stage variant-specific vaccine. Moreover, the immune responses, specificity, cross-reactivity, and host cell gene expression landscapes upon vaccination have to be rigorously tested for such variant-specific vaccines to be developed. To directly assess the immunogenicity of potential variant-specific SARS-CoV-2 vaccination, we generated LNP-mRNA vaccine candidates that encode the B.1.351 and B.1.617 spikes, along with the WT spike. With these variant-specific LNP-mRNAs, we characterized the immune responses they induce in animals against homologous (cognate) and heterologous spike antigens and SARS-CoV-2 pseudoviruses. To better understand the systematic immune responses induced by variant-specific SARS-CoV-2 spike mRNA-LNP vaccination, we analyzed the combined single-cell transcriptomes and lymphocyte antigen receptor repertoires of mice immunized with B.1.351 and B.1.617 spike mRNA-LNP vaccine candidates.

## Results

### Design, generation and physical characterization of variant-specific SARS-CoV-2 spike LNP-mRNAs

We designed and generated nucleotide-modified mRNAs separately encoding full-length SARS-CoV-2 WT, B.1.351 and B.1.617 spikes. The HexaPro mutations(Wrapp et al., 2020) were introduced and the furin cleavage site(Laczko et al., 2020) was replaced with a GSAS sequence to stabilize the prefusion state and preserve integrity of spike S1 and S2 subunits (Figure 1a-1b). The protein expression and receptor binding ability of modified spike mRNA were confirmed by *in vitro* cell transfection and flow cytometry where the spike binding to the human ACE2-Fc fusion protein was detected by PE-anti-Fc antibody (Figure S 1a-b). We encapsulated the spike mRNA with LNP, of which size and homogeneity were evaluated by dynamic light scattering (DLS) and transmission electron microscopy (TEM). The WT, B.1.351 and B.1.617 mRNA LNPs have mean diameters of 80.7 ± 6.9nm, 66.4 ± 5.3 nm and 72.2 ± 5.8 nm with a monodispersed size distribution as determined by DLS and polydispersity indices of 0.08, 0.13 and 0.08, respectively (Figure S 1c-1d). The immunogenicity of the LNP-mRNA was assessed in C57BL/6Ncr mice by two intramuscular injections (doses) of 1 µg or 10 µg LNP-encapsulated mRNA, separated by 3 weeks (prime and boost, respectively) (Figure 1c). Serum samples were collected two weeks after the prime and boost, and then subjected to ELISA and neutralization assays to evaluate the antibody response. These mice were sacrificed 40 days post vaccination, and the spleen, lymph nodes and blood cells were collected for downstream assays, including single cell transcriptomics sequencing (scRNA-seq), bulk and single cell BCR sequencing (BCR-seq) and TCR sequencing (TCR-seq), as well as flow cytometry. All procedures were standardized across all groups.

**Figure 1.**
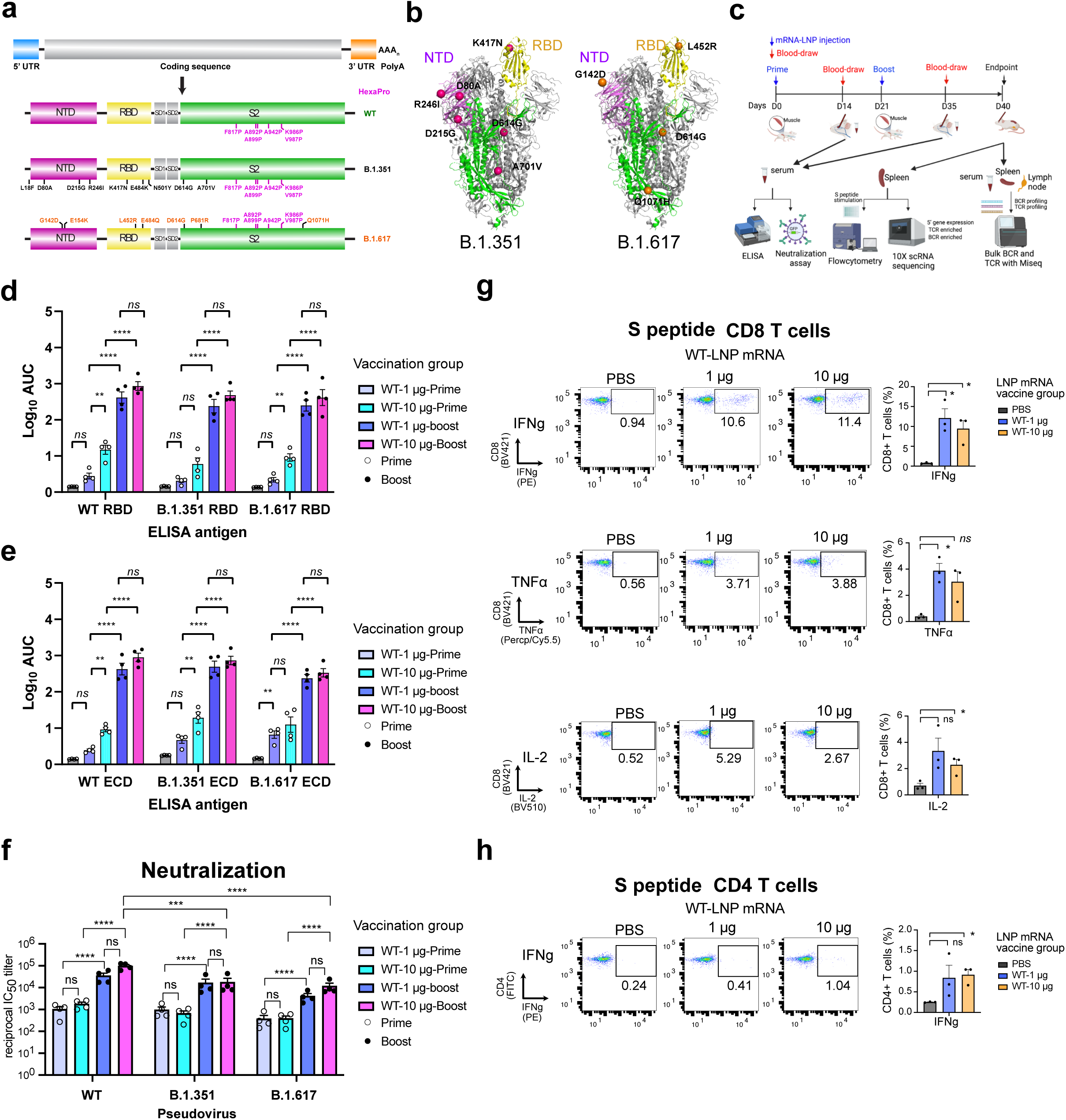
Overview of the primary experimental design, and the B and T cell responses induced by WT-LNP-mRNA vaccination against SARS-CoV-2 WT, B.1.351 and B.1.617 spikes in mice. **a**, Schematic of the designs of three variant-specific LNP-mRNA vaccine candidates. Functional elements were shown in the spike mRNA and translated protein of SARS-CoV-2 WT, B.1.351 and B.1.617 spikes, including protein domains, HexaPro and variant-specific mutations. **b**, 3D structure highlighting certain variant-specific mutations in B.1.351 and B.1.617 spikes. Distribution of mutations of B.1.351 and B.1.617 were shown in the structure of SARS-CoV-2 (PDB: 6VSB). Mutations of B.1.351 and B.1.617 were shown as spheres, except for those in the unstructured loop regions. Certain mutations were not visible in the structure as they fall into floppy regions of spike. **c**, Schematic of overall design of primary experiments. Six- to 8-week-old C57BL/6Ncr mice (B.1.351-LNP-mRNA (top) and B.1.617-LNP-mRNA, n = 6 mice per group; WT-LNP-mRNA, n = 4 mice; PBS, n = 9) received 1 or 10 µg of WT-LNP mRNA, B.1.351-LNP-mRNA or B.1.617-LNP-mRNA via the intramuscular route on day 0 (Prime) and day 21 (Boost). Blood was collected twice, two weeks post prime and boost. The binding and pseudovirus-neutralizing antibody responses induced by LNP-mRNA were evaluated by ELISA and neutralization assay. Mice were euthanized at day 40. The spleen, lymph node and blood samples were collected to analyze immune responses in by flow cytometry, bulk BCR and TCR profiling and single cell profiling. **d-e,** Serum ELISA titers of WT-LNP mRNA vaccinated animals (n = 4). Serum antibody titer as area under curve (AUC) of log_10_-transformed curve (1og_10_ AUC) to spike RBDs (**d**) and ECDs (**e**) of SARS-CoV-2 WT, B.1.351 and B.1.617. Two-way ANOVA with Tukey’s multiple comparisons test was used to assess statistical significance. **f,** Serum neutralization titers of WT-LNP mRNA vaccinated animals (n = 4). Cross neutralization of SARS-CoV-2 WT, B.1.351 or B.1.617 pseudovirus infection of ACE2-overexpressed 293T cells. Two-way ANOVA with Tukey’s multiple comparisons test was used to assess statistical significance. **g-h**, T cell response of WT-LNP mRNA vaccinated animals (n = 4). CD8^+^ (**g**) and CD4^+^ (**h**) T cell responses were measured by intracellular cytokine staining 6 hours after addition of BFA. The unpaired parametric t test was used to evaluate the statistical significance. **Notes:** In this figure: Each dot represents data from one mouse. Data are shown as mean ± s.e.m. plus individual data points in dot plots. Statistical significance labels: n.s., not significant; * p < 0.05; ** p < 0.01; *** p < 0.001; **** p < 0.0001 Source data and additional statistics for experiments are provided in a supplemental excel file. **See also: Figure(s) S1, S2**

### Immune responses induced by WT-LNP-mRNA vaccination in mice

WT-LNP-mRNA induced dose-dependent binding antibody responses against spike ECD and RBD of SARS-CoV-2 WT, B.1.351 and B.1.617 variants after prime and boost (Figure 1d-e). Compared to the post-prime immune response, orders of magnitudes increase in immune response were observed after the boost injection, suggesting that the second dose significantly boosted B cell immunity to SARS-CoV-2 antigen (Figure 1d-e). Using a pseudovirus neutralization assay that has been widely reported to be consistent with authentic virus results (Chen et al., 2021; Schmidt et al., 2020a), the serum samples from mice receiving WT-LNP-mRNA vaccination also showed potent neutralization activity against all three variants, again with a strong prime-boost effect (Figure 1f). However, the neutralization ability of WT-LNP-mRNA vaccinated sera was found to be several fold lower against either B.1.351 or B.1.617 as compared to the cognate WT pseudovirus (Figure 1f). These observations are consistent with the series of reports showing dramatic reduction in neutralization of B.1.351 and B.1.617 variants by vaccinated individuals’ sera, convalescent sera, and therapeutic antibodies (Edara et al., 2021; Planas et al., 2021a; Wang et al., 2021b; Zhou et al., 2021a).

To evaluate the T cell response to the spike peptides, the splenocytes were isolated from mouse spleens 40 days post vaccination and the antigen-specific CD4^+^ and CD8^+^ T cell response to S peptide pools were determined by intracellular cytokine staining (Figure S 2). WT-LNP-mRNA, at both low and high doses, induced reactive CD8^+^ T cells producing interferon γ (IFN-γ, IFNg), tumor necrosis factor *α* (TNF-*α*, TNFa), and interleukin 2 (IL-2) (Figure 1g), at levels consistent with previously reported studies(Corbett et al., 2020; Laczko et al., 2020). WT-LNP-mRNA at both doses also induced reactive CD4^+^ T cells that produce IFN-γ^+^, but little for TNF-*α*, IL-2, IL-4 or IL-5 (Figure S 2b). As technical quality controls, there is no difference in cytokine production between vaccinated groups and the PBS group when cells were treated with vehicle (no peptide) or PMA/ionomycin (Figure S 2c-d). These results suggest that WT-LNP-mRNA vaccines are able to induce potent spike protein specific CD4 and CD8 T cell responses.

### Binding and neutralizing antibody responses of B.1.617-LNP-mRNA and B.1.351-LNP-mRNA

Both B.1.617-LNP-mRNA and B.1.351-LNP-mRNA induced dose-dependent binding antibody responses against spike ECD and RBD of SARS-CoV-2 WT, B.1.351 and B.1.617 variants (Figure 2a-2b; Figure S 3). The strong boost effect in ELISA was also observed for these two variant-specific LNP-mRNAs (Figure 2a-2b; Figure S 3a-b). The dose-dependence effect was observed in both B.1.617-LNP-mRNA and B.1.351-LNP-mRNA groups across three types of ELISA antigens of both RBD and ECD, although the dose effect was less prominent in the post-boost samples, where both doses showed high titers at potential saturation level (Figure 2a-2b; Figure S 1e-f). Relatively speaking, higher antibody responses was often observed with ECD antigen, suggesting an immunogenic domain other than RBD contributed to the additional response to spike ECD (Figure 2a-2b; Figure S 1e-f). Overall, the binding intensity as measured by serum titer between RBD and ECD strongly correlates with each other across all vaccination groups (Figure S 1g).

**Figure 2.**
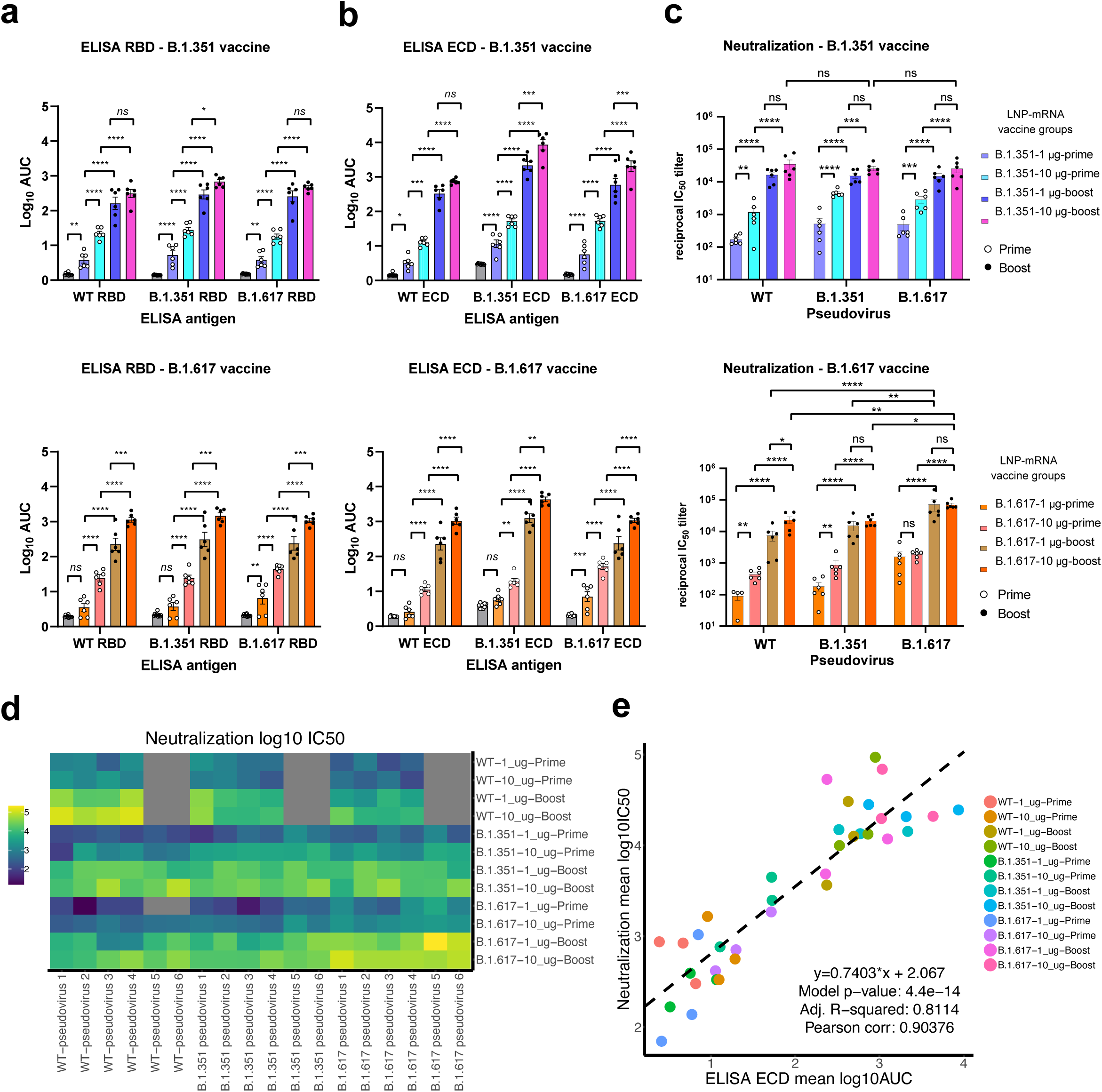
B.1.351-LNP-mRNA and B.1.617-LNP-mRNA elicit robust binding and pseudovirus-neutralizing antibody response against all three variants in mice. **a,** Serum ELISA titers of animals vaccinated by B.1.351-LNP-mRNA (top) and B.1.617-LNP-mRNA (bottom), against RBD from three different spikes (WT, B.1.351, and B.1.617) of SARS-CoV-2 (n = 6). **Notes:** In this figure: Each dot represents data from one mouse. Data are shown as mean ± s.e.m. plus individual data points in dot plots. Statistical significance labels: n.s., not significant; * p < 0.05; ** p < 0.01; *** p < 0.001; **** p < 0.0001 Source data and additional statistics for experiments are provided in a supplemental excel file. **See also: Figure(s) S1**

We then examined pseudovirus-neutralizing antibody response. Both B.1.617-LNP-mRNA and B.1.351-LNP-mRNA elicited potent neutralizing antibodies, of which response mirrored the trend of post-prime and post-boost response reported in ELISA (Figure 2c-2d). The initial level of neutralization was at 10^2^ – 10^3^ level of reciprocal IC50 after priming for most groups (Figure 2c-2d). Consistent with findings in ELISA, an approximately two orders of magnitude increase in neutralization titer by boost was observed across all groups (for both vaccine candidates and for all three pseudovirus types) in the low dose (1 µg) setting, and there was an approximately one order of magnitude increase in the high dose (10 µg) setting (Figure 2c-2d). The dose effect of serum neutralization activity for both B.1.617-LNP-mRNA and B.1.351-LNP-mRNA was observed at priming for most groups, but negligible post boost (i.e. both 1 µg and 10 µg dose groups reach reciprocal IC50 titer of 10^4^ level after boost) (Figure 2c-2d). Both B.1.617-LNP-mRNA and B.1.351-LNP-mRNA effectively neutralize pseudoviruses of all three SARS-CoV-2 pseudoviruses post boost at titers of 10^4^ level (Figure 2c-2d). Interestingly, B.1.351-LNP-mRNA vaccinated animals neutralize pseudoviruses of all three SARS-CoV-2 at similar levels post boost at both doses (Figure 2c-2d); while B.1.617-LNP-mRNA vaccinated animals showed significantly higher titer against its cognate B.1.617 pseudovirus (by several folds). Overall, across all vaccination groups, the neutralization activity strongly correlates with binding intensity for ECD binding (Figure 2e), which also holds true for RBD binding (Figure S 1h).

### B.1.617-LNP-mRNA and B.1.351-LNP-mRNA elicited strong systemic T cell response against SARS-CoV-2 spike

Similarly, to evaluate the T cell response to the spike peptides, the splenocytes were isolated from mouse spleens 40 days post vaccination and the antigen-specific CD4^+^ and CD8^+^ T cell responses to S peptide pools were determined by intracellular cytokine staining (Figure S 2e-g). Positive control PMA/ionomycin treatment group and negative control no peptide group were both validated (Figure S 2f-g). Both B.1.617-LNP-mRNA and B.1.351-LNP-mRNA, at low and high doses, induced potent reactive CD8^+^ T cell responses in terms of cellular production of IFN-γ, TNF-*α*, and IL-2 (Figure 3a-3c). Both LNP-mRNAs at both doses also induced reactive CD4^+^ T cells that produce IFN-γ, but only minimally for TNF-*α*, and had no effect on IL-2, IL-4 or IL-5 (Figure S 2e).

**Figure 3.**
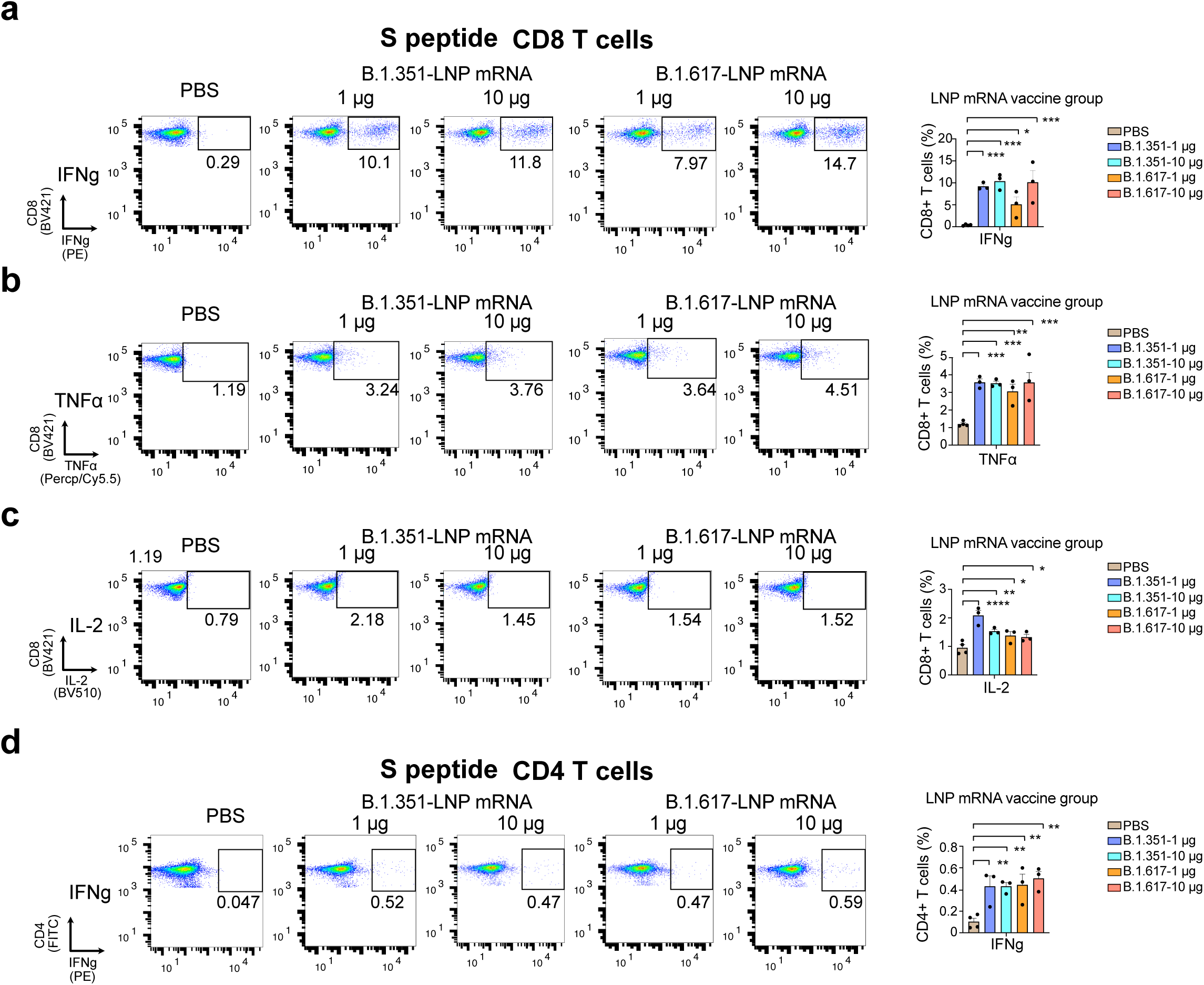
B.1.351-LNP-mRNA and B.1.617-LNP-mRNA induced S protein-specific T cell response. **a-c** Percentage of CD8^+^ T cells expressing IFN-γ (**a**), TNFα (**b**), and IL-2 (**c**), in response to stimulation of S peptide pools (n=3). Left panels, representative flow plots; right panels, dot-bar plots for statistics of the left panels. **d**, Percentage of CD4^+^ T cells expressing IFN-γ in response to stimulation of S peptide pools (n = 3). Left panels, representative flow plots; right panels, dot-bar plots for statistics of the left panels. **Notes:** In this figure: Each dot represents data from one mouse. Data are shown as mean ± s.e.m. plus individual data points in dot plots. Statistical significance labels: n.s., not significant; * p < 0.05; ** p < 0.01; *** p < 0.001; **** p < 0.0001 Source data and additional statistics for experiments are provided in a supplemental excel file. **See also: Figure(s) S2**

### Single cell immune repertoire mapping of B.1.617-LNP-mRNA and B.1.351-LNP-mRNA vaccinated animals

In order to gain insights on the global composition and transcriptional landscape of the immune cells, we performed single cell transcriptomics (scRNA-seq) on the spleen samples of B.1.351-LNP-mRNA and B.1.617-LNP-mRNA vaccinated animals. As visualized in an overall UMAP, from a total of 16 animals from 4 vaccination groups (B.1.351-LNP-mRNA and B.1.617-LNP-mRNA, both 1 µg and 10 µg dose groups), plus a control group (PBS treated), we sequenced the transcriptomes of a total of 90,152 single cells, as projected on a Uniform Manifold Approximation and Projection (UMAP) (Figure 4a; Figure S 3a). Clustering was performed with Louvain algorithm, which identified 21 clusters from respective signatures of their differentially expressed genes (Figure 4b; Figure S 3b). With the expression of a number of cell type specific markers, such as markers for pan-leukocytes (*Ptprc/Cd45*), B cells (*Cd19, Cd22*), plasma cells (*Sdc1/Cd138*), T cells (*Cd3e, Cd4, Cd8a, Cd8b1, Trac/TCRa*), natural killer (NK) cells (*Ncr1, Klrb1c*), dendritic cells / macrophages / monocytes (*Cd11b/Itgam, Cd11c/Itgax, Adgre/F4/80, Mrc1, Gsr1*), red blood cells (RBCs) (*Hba-a1*), and neutrophils (*S100a8, Mmp9*), we assigned cellular identities to the clusters, which include B cells (*Cd19^+^*), progenitor B cells (*Csf1r^+^;Cd19^+^*), plasma cells (*Igha^+^/Ighm^+^;Sdc1^+^;Cd19^-^*), B cell-like cells (*Cd19^+^;Ly6a^+^;*), CD4 T cells (*Cd3e^+^;Cd4^+^*), CD8 T cells (*Cd3e^+^;Cd8a^+^;Cd8b1^+^*), NKT cells (*Klra1^+^;Klra6^+^;Zbtb16^+^*), DCs (*Itgam^+^;Itgax^+^*), macrophages (*Itgam^+^;Csf1r^+^;Adgre^+^;Mrc1^+^*), monocytes (*Itgam^+^;Csf1r^+^;Gsr1^+^*), neutrophils (*S100a8^+^/S100a9^+^;Mmp9^+^*), NK cells (*Cd3e^-^;Ncr1^+^,Klrb1c^+^*), and RBCs (*Ptprc^-^;Hba-a1^+^*) (Figure 4a, 4c; Figure S 3c). Noted that while Cluster 5 contains predominantly CD8 T cells, it also contains a small population of NKT cells that were not separated by the automatic clustering algorithm (Figure 4a; Figure S 3d). The single cell transcriptomics provided a landscape of systemic immune cell populations and their respective gene expression (GEX) data in B.1.351-LNP-mRNA and B.1.617-LNP-mRNA vaccinated along with placebo control animals.

**Figure 4.**
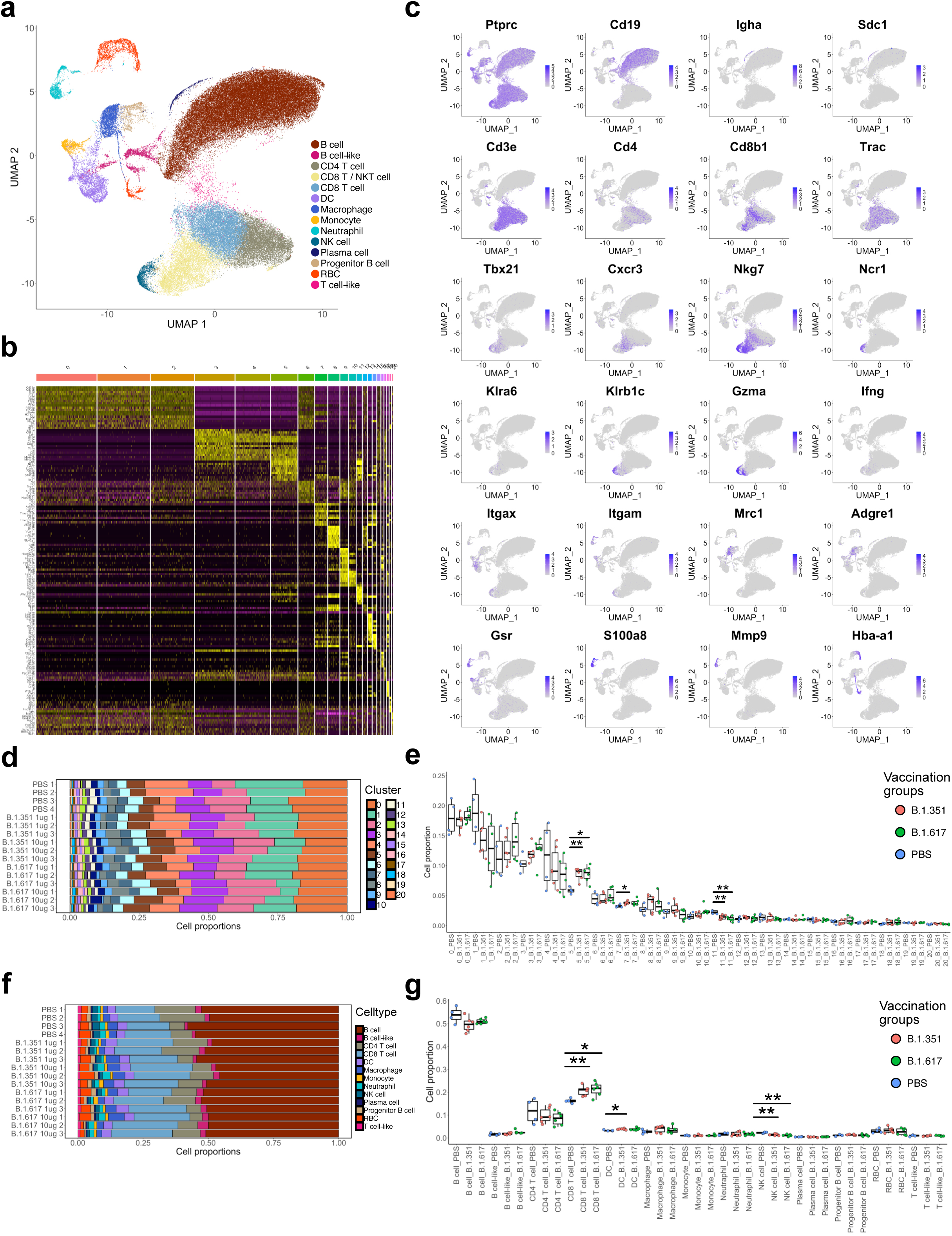
Single cell transcriptomics and immune repertoire profiling of B.1.351-LNP-mRNA and B.1.617-LNP-mRNA vaccinated animals. **a,** Overall UMAP, visualization of all 90,152 cells pooled across samples and conditions. Clustering was performed with Louvain algorithm with 21 identified clusters, and assignment of cell identity was performed using expression of cell type specific markers. **b,** Heatmap of differentially expressed markers for all clusters. **c,** Overall UMAPs, with expression of a number of cell type specific markers. **d,** Bar chart depicting the cell proportion of clusters for each sample. **e,** Boxplots of cell proportions by clusters for each condition (PBS, n = 4; B.1.351-LNP-mRNA, n = 6; B.1.617-LNP-mRNA, n = 6). Comparison between groups performed with Wilcoxon ranked sum test. **f,** Bar chart depicting the cell proportions of different immune cell types for each sample. **g,** Boxplots of cell proportions by cell type for each condition (PBS, n = 4; B.1.351-LNP-mRNA, n = 6; B.1.617-LNP-mRNA, n = 6). Comparison between groups performed with Wilcoxon ranked sum test. **Notes:** In this figure, panels **e** and **g**: Each dot represents data from one mouse. The high dose (n = 3 each) and low dose (n = 3 each) groups for each vaccine were merged (n = 6 total) in single cell data analysis, same thereafter. Statistical significance labels: n.s., not significant; * p < 0.05; ** p < 0.01; *** p < 0.001; **** p < 0.0001 **See also: Figure(s) S3**

We then compared the systemic (spleen) immune cell compositions between placebo and vaccinated animals (Figure 4d-g). Out of the 21 clusters, three have significantly changed fractions in the total splenocytes upon vaccination as compared to placebo, including a significant increase in Cluster 5 (composed of CD8 T cells and NKT cells) for both B.1.351-LNP-mRNA and B.1.617-LNP-mRNA, a slight increase in Cluster 7 (DCs) for B.1.351-LNP-mRNA, and a slight decrease in Cluster 11 (NK cells) for both LNP-mRNAs (Figure 4d-e). Analysis by summing all of the same cell types from different clusters validated this finding, revealing a significant increase in CD8 T cells (after excluding NKT cells) for both for B.1.351-LNP-mRNA and B.1.617-LNP-mRNA, a slight increase in DCs for B.1.351-LNP-mRNA, and a slight decrease in NK cells for both vaccination groups (Figure 4d-e). We noted that a fraction of Cluster 5’s CD8 T cells are also marked by the positive expression of *Nkg7*, *Ifng, Gzma,* and *Cxcr3*, representing a cluster of more activated T cells (Figure 4b-c; Figure S 3c). The increase in CD8 T cells, especially the cluster of more activated T cells, is consistent with the strong CD8 T cell responses as detected by flow cytometry (Figure 3a-c), indicative of systemic CD8 T cell responses upon vaccination with these variant-specific LNP-mRNAs.

### Gene expression signatures of B cell and T cell populations of B.1.617-LNP-mRNA and B.1.351-LNP-mRNA vaccinated animals

Because B and T cells are the cornerstones of adaptive immunity against SARS-CoV-2, we further investigated the B cell sub-populations and T cell sub-populations, respectively. Using the global clustering results with a number of B cell lineage markers (Figure 4a; Figure 5a; Figures S 5-6), we identified a total of 49,236 B cell-associated populations from all samples and conditions of the 16 mice (Figure 5b). Using unbiased clustering, these B sub-population cells are divided into 15 Clusters (Figure 5b; Figure S 4a-b), although the largest 8 clusters are near each other in UMAP space and form a “supercluster”. Similarly, using the global clustering results with T cell lineage markers (Figure 4a; Figure 5b; Figures S 5 and 7), we identified a total of 28,099 T cell-associated populations (Figure 5d). Using unbiased clustering, these T sub-population cells are divided into 12 Clusters (Figure 5d; Figure S 4c-d). Using more refined T cell markers, cells that represent sub-classes of T cells can be detected, such as CD4 T cells (*Cd3e^+^;Cd4^+^*), CD8 T cells (*Cd3e^+^;Cd8a^+^;Cd8b1^+^*), regulatory T cells (Tregs) (*Cd3e^+^;Cd4^+^;Foxp3*^+^), Th1-like T helper cells (Th1s) (*Cd3e^+^;Cd4^+^;Cxcr6^+^;Tbx21/Tbet^+^;Stat4*^+^), Th2-like T helper cells (Th2s) (*Cd3e^+^;Cd4^+^;Ccr4^+^;Il4ra^+^;Stat6*^+^), Th17-like T helper cells (Th17s) (Tregs) (*Cd3e^+^;Cd4^+^;Rorc^+^;Stat3*^+^), and T follicular helper cells (Tfhs) (*Cd3e^+^;Cd4^+^;Cd40lg^+^;Il4^+^;Il21*^+^) (Figure S 4c-d). Noted that as observed in various single cell datasets (Lindeboom et al., 2021), gating cellular populations by gene expression of these markers do not always produce clear cut populations defined by canonical immune markers using flow cytometry, possibly due to the differences between mRNA vs. surface protein expression, as well as the pleiotropic roles of various genes.

**Figure 5.**
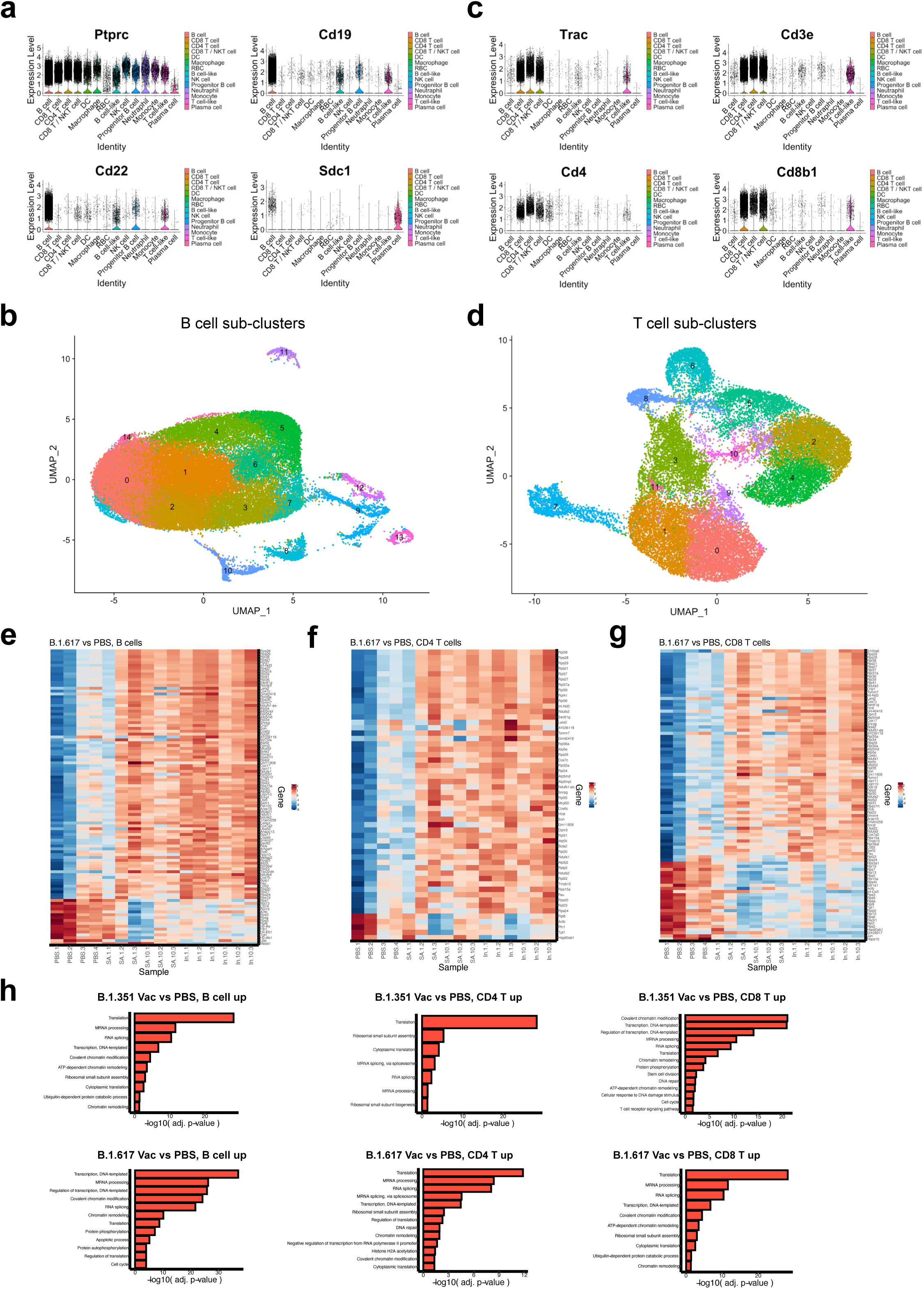
Single cell analysis of B cell and T cell populations and gene expression signatures of B.1.351-LNP-mRNA and B.1.617-LNP-mRNA vaccinated animals. **a,** Violin plots of expression of a number of cell type specific markers that define B cell and plasma cell lineages. **b,** B cell UMAP, visualization of 49,236 B cell-associated populations pooled across samples and conditions. Clustering was performed with Louvain algorithm with 15 identified clusters. **c,** Violin plots of expression of a number of cell type specific markers that define T cell lineages. **d,** T cell UMAP, visualization of 28,099 T cell-associated populations pooled across samples and conditions. Clustering was performed with Louvain algorithm with 12 identified clusters. **e,** Heatmap of differentially expressed genes between B.1.617-LNP-mRNA sample B cells vs PBS sample B cells, expression profiles shown for all samples (n = 16). **f,** Heatmaps of differentially expressed genes between B.1.617-LNP-mRNA sample CD4 T cells vs PBS sample CD4 T cells and B.1.617-LNP-mRNA sample CD4 T cells vs PBS sample CD4 T cells. Expression profiles shown for all samples (n = 16). **g,** Heatmaps of differentially expressed genes between B.1.617-LNP-mRNA sample CD8 T cells vs PBS sample CD8 T cells and B.1.617-LNP-mRNA sample CD8 T cells vs PBS sample CD8 T cells. Expression profiles shown for all samples (n = 16). **h,** Bar charts depicting significance values for enriched Gene Ontology Biological Process terms associated with upregulated genes for B.1.351-LNP-mRNA or B.1.617-LNP-mRNA vs PBS group, in B cells, CD4 T cells or CD8 T cells. **See also: Figure(s) S4, S5**

To examine the transcriptomic changes in the B and T cell sub-populations upon vaccination, we performed differential expression (DE) analysis in the matched sub-populations between PBS and B.1.351-LNP-mRNA, or B.1.617-LNP-mRNA, groups. Vaccination caused differential expression of genes in host B cells, CD4 T cells and CD8 T cells (Figure 5e-g; Figures S 5). The differentially expressed genes intersect with genes in B cell activation, immune effector, and immune response genes, such as *Lyn, Cd22* and *Btla* (Figure S 5a-b). The differentially expressed genes in CD4 and CD8 T cells also intersect with genes in T cell activation, immune effector, and immune response genes, such as *Cd40lg, Perforin/Prf1, Dhx36, Ddx17, Ddx21, Ccl5, Il18r1, Ptpn22* and *Plcg1* (Figure S 5c-e). Interestingly, the top upregulated expressed genes in B cells represent transcription and translation machineries, which is consistent between B.1.351-LNP-mRNA and B.1.617-LNP-mRNA vaccination groups (Figure 5h; Figure S 5a-b). This strong signature was also observed in T cells (Figure 5h; Figure S 5c-e), consistent with the phenomenon of active lymphocyte activation upon vaccination (Clem, 2011; Jeyanathan et al., 2020; Teijaro and Farber, 2021).

### TCR and BCR diversity mapping of B.1.617-LNP-mRNA and B.1.351-LNP-mRNA vaccinated animals

To reveal the B and T cell clonal diversity and influence by vaccination, we performed VDJ repertoire mapping and clonal analyses of B cell and T cell populations of B.1.351-LNP-mRNA and B.1.617-LNP-mRNA vaccinated animals. We performed both single cell BCR sequencing (scBCR-seq) and single cell TCR sequencing (scTCR-seq) on the spleen samples of all groups (4 vaccination groups and a PBS group, n = 16 mice total). We sequenced a total of 47,463 single B cells and 25,228 single T cells. Clonal composition showed the BCR repertoire in the single cell BCR-seq dataset, revealing a trend towards decreased clonal diversity (Figure 6a-b; Figure S 6a-b). The clonal composition of single cell TCR showed a significant decrease in clonal diversity (Figure 6c-d; Figure S 6c-d). This phenomenon is consistent with the clonal expansion of stimulated lymphocytes upon vaccination, which is more pronounced in the scTCR-seq data.

**Figure 6.**
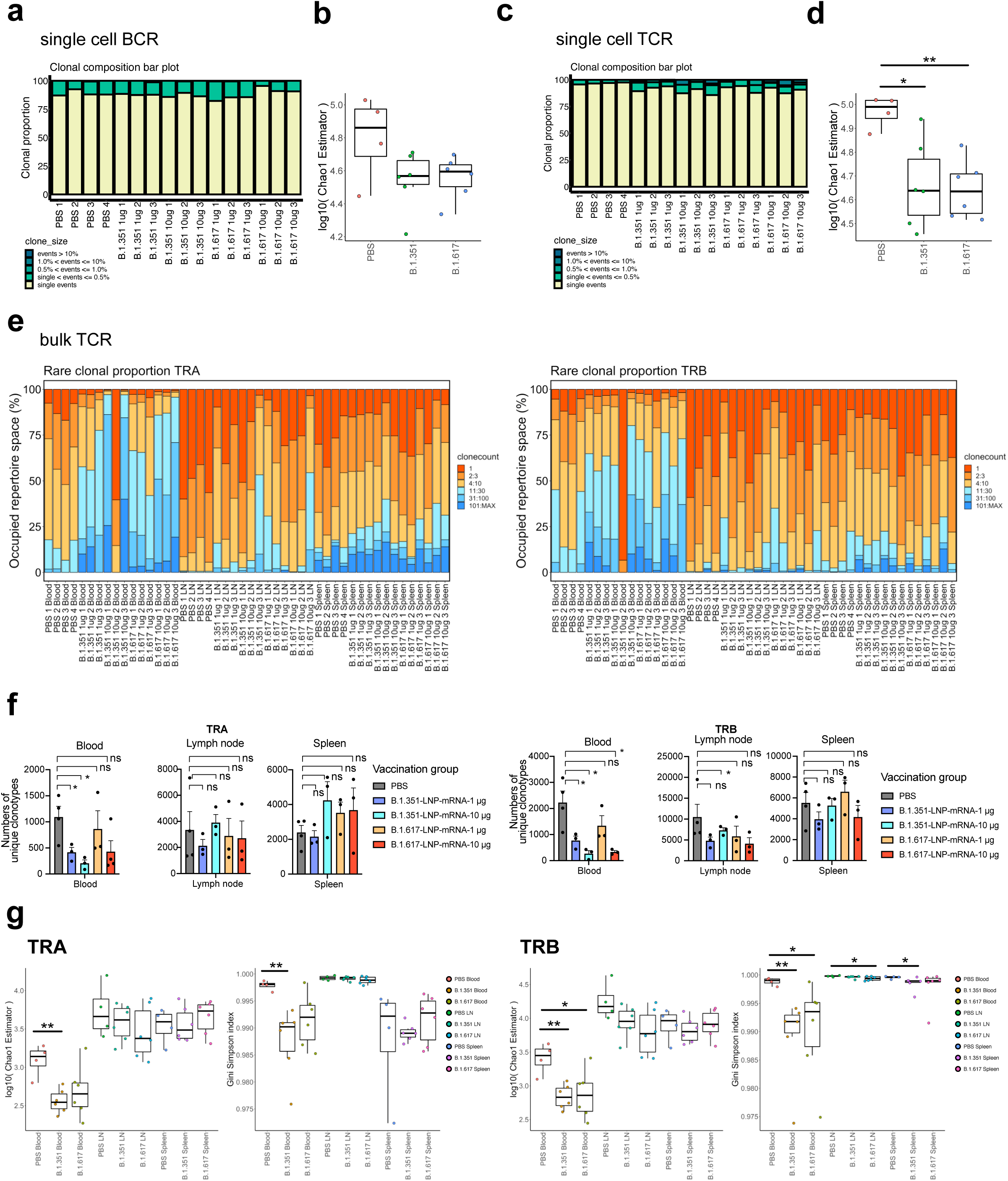
VDJ repertoire and clonal analyses of B cell and T cell populations of B.1.351-LNP-mRNA and B.1.617-LNP-mRNA vaccinated animals. **a,** Clonal composition bar plot depicting proportion of the BCR repertoire occupied by the clones of a given size for all samples (n = 16) in the single cell BCR-seq dataset. **b,** Boxplot of Chao1 indices for each condition (PBS, n = 4; B.1.351-LNP-mRNA, n = 6; B.1.617-LNP-mRNA, n = 6) for repertoires in the single cell BCR-seq dataset. **c,** Clonal composition bar plot depicting proportion of the TCR repertoire occupied by the clones of a given size for all samples (n = 16) in the single cell TCR-seq dataset. **d,** Boxplot of Chao1 indices for each condition (PBS, n = 4; B.1.351-LNP-mRNA, n = 6; B.1.617-LNP-mRNA, n = 6) for repertoires in the single cell TCR-seq dataset. **e,** Bar plot depicting relative abundance for groups of rare clonotypes for TRA and TRB chains for all samples (n = 48) in the bulk TCR-seq dataset. **f,** Bar plot depicting number of unique clonotypes for TRA and TRB chains in the bulk TCR-seq dataset across vaccination and tissue of origin groups. **g,** Boxplots of Chao1 (left) and Gini-Simpson (right) indices for TRA or TRB chain repertoires in the bulk TCR-seq dataset across vaccination and tissue of origin groups. **Notes:** In this figure, panels **b, d, f** and **g**: Each dot represents data from one mouse. The low dose and high dose groups of the same vaccine were grouped together. Statistical significance labels: n.s., not significant; * p < 0.05; ** p < 0.01; *** p < 0.001; **** p < 0.0001 **See also: Figure(s) S6, S7**

To further validate the observations, we also performed bulk BCR-seq and bulk TCR-seq for all these mice on additional tissue samples, i.e. spleen, peripheral blood (PB) and lymph node (LN). The bulk BCR-seq and TCR-seq data revealed systematic clonality maps of the spleen, PB and LN samples of the variant-specific LNP-mRNA vaccinated along with placebo treated animals (Figure 6e-g; Figure S 7a-b). The bulk TCR-seq results, from both TCR alpha and beta chains (*TRA* and *TRB*), validated the observation of decreased clonal diversity from single cell VDJ profiling (Figure 6e-g), again consistent with the notion of clonal expansion of a small number of clones. Interestingly, the clonal diversity decrease / clonal expansion effect is strongest in the PB samples that capture circulating T cells (Figure 6e-g). These data together unveiled BCR and TCR repertoire clonality, diversity and respective shifts in variant-specific LNP-mRNA vaccinated animals as compared to placebo-treated.

## Discussion

Although efficacious COVID-19 mRNA vaccines have been deployed globally, the rapid spread of SARS-CoV-2 VoCs with higher contagiousness as well as resistance to therapies and vaccines demands evaluation of next-generation COVID-19 vaccines specifically targeting these evolving VoCs. Mounting evidence has suggested that the B.1.351 and B.1.617 lineage variants of SARS-CoV-2 possesses much stronger immune escape capability than the original wildtype virus (Planas et al., 2021a; Wang et al., 2021b). The Delta variant has swept over the population and became the dominant variant in late 2021 (https://www.gisaid.org). More recently, a heavily mutated Omicron variant (B.1.1.529) quickly arose in frequency in South Africa (Callaway, 2021b) (Pulliam et al. 2021), and spread across the globe (https://www.gisaid.org). The lower neutralizing titers in fully vaccinated patients were found associated with breakthrough infections (Bergwerk et al., 2021). It has been speculated that the waning immunity from early vaccination and emergence of more virulent SARS-CoV-2 variants may lead to reduction in vaccine protection and increase of breakthrough infections(Callaway, 2021a; Pegu et al., 2021). It has been reported that mRNA vaccines’ efficacy against B.1.351 and B.1.617.2 dropped significantly(Abu-Raddad et al., 2021; Lopez Bernal et al., 2021b). Moreover, for individuals receiving only a single dose of vaccine, the protective efficacy can be dramatically lower (Iacobucci, 2021). It is worth noting that efficacy value and definition may vary from study to study(Charmet et al., 2021; Planas et al., 2021b), which were conducted in different regions and populations. All these factors urged for next-generation variant-specific COVID-19 vaccines, and prompted us to evaluate the mRNA vaccine candidates encoding VoC spikes as antigens.

Our study characterized the titers and cross-reactivity of sera from mice vaccinated with WT-, B.1.351- or B.1.617-LNP-mRNAs to all three WT, B.1.351 and B.1.617 spike antigens and pseudoviruses. In agreement with findings in patients’ sera, we found that the neutralizing titers of WT vaccine sera were several folds lower against the two variants of concerns than against WT pseudovirus. Interestingly, the B.1.617-LNP-mRNA vaccinated sera also showed particularly strong neutralization activity against its cognate B.1.617 pseudovirus, while the B.1.351-LNP-mRNA showed similar neutralization activity against all three pseudoviruses. It is worth noting that all three forms of vaccine candidates can induce potent B and T cell responses to WT as well as the two VoCs’ spikes.

The T cell-biased immune response is important for antiviral immunity and thereby the efficacy and safety of viral vaccines (Graham, 2020). To evaluate the Th1 and Th2 immune response by the variant vaccines, we performed intracellular staining of Th1 and Th2 cytokines in splenocytes. After stimulation with peptide pools covering the entire S protein, the splenocytes from three mRNA vaccine groups produced more hallmark Th1 cytokine IFN-γ in both CD4^+^ and CD8^+^ T cells than those from PBS group. Our flow cytometry data suggested that the two variant vaccine candidates induced strong Th1-biased immune responses, just like the WT vaccine, of which Th1 response had been observed by previous studies in animal models (Corbett et al., 2020; Laczko et al., 2020).

Single cell sequencing is a powerful technology for immune and gene expression profiling, which has been utilized for mapping immune responses to COVID-19 infection (Stephenson et al., 2021; Zhang et al., 2020). In order to gain insights on the transcriptional landscape of the immune cells, and clonal repertoire changes specifically in B and T cells, we performed single cell transcriptomics, as well as BCR and TCR repertoire sequencing. The single cell transcriptomics data revealed a systematic landscape of immune cell populations in B.1.351-LNP-mRNA and B.1.617-LNP-mRNA vaccinated animals. We mapped out the repertoires and associated global gene expression status of the immune populations including B cells, T cells, and innate immune cells. From the overall splenocyte population, we observed a distinct and significant increase in the CD8 T cell populations in vaccinated animals. Interestingly, differential expression between vaccinated and placebo-treated animals showed a strong signature of increased expression of transcriptional and translational machinery in both B and T cells. While the actual mechanism awaits future studies, these phenomena are potentially reflective of the active proliferation and immune responses in these lymphocytes.

BCR and TCR sequencing are efficient tools for mapping of clonal repertoire diversity, which has been rapidly utilized for sequencing COVID-19 patients (Schultheiss et al., 2020; Zhang et al., 2020). BCR-seq and TCR-seq unveiled the diversity and clonality and respective shifts in variant-specific LNP-mRNA inoculated animals as compared to placebo-treated. The decrease in VDJ clonal diversity, along with clonal expansion of a small number of clones, are observed in vaccinated animals as compared to placebo group. Vaccinated animals from both B.1.351-LNP-mRNA and B.1.617-LNP-mRNA groups have clonal TCR expansion, especially pronounced in peripheral blood samples. The induction of diverse and expanding clones is a signature of vaccine induced protective immunity (Clem, 2011).

Our study provided direct assessment of *in vivo* immune responses to vaccination using LNP-mRNAs encoding specific SARS-CoV-2 variant spikes in pre-clinical animal models. The single cell and bulk VDJ repertoire mapping also provided unbiased datasets and robust systems immunology of SARS-CoV-2 vaccination by LNP-mRNA specifically encoding B.1.351 and B.1.617 spikes. These original data may offer valuable insights for the development of the next-generation COVID-19 vaccines against the SARS-CoV-2 pathogen and especially its emerging variants of concern (Singh et al., 2021). The efficacy and safety of the variant-specific vaccine candidates need to be rigorously tested in future translational and clinical studies.

## Methods

### Plasmid construction

The DNA sequences of B.1.351 and B.1.617 SARS-CoV-2 spikes for the mRNA transcription and pseudovirus assay were synthesized as gBlocks (IDT) and cloned by Gibson Assembly (NEB) into pcDNA3.1 plasmids. To improve expression and retain prefusion conformation, six prolines (HexaPro variant, 6P) were introduced to the SARS-CoV-2 spike sequence in the mRNA transcription plasmids. The plasmids for the pseudotyped virus assay including pHIV_NL_GagPol and pCCNanoLuc2AEGFP are gifts from Dr. Bieniasz’ lab(Schmidt et al., 2020b). The C-terminal 19 amino acids were deleted in the SARS-CoV-2 spike sequence for the pseudovirus assay.

### Cell culture

HEK293T (ThermoFisher) and 293T-hACE2 (gifted from Dr Bieniasz’ lab) cell lines were cultured in complete growth medium, Dulbecco’s modified Eagle’s medium (DMEM; Thermo fisher) supplemented with 10% Fetal bovine serum (FBS, Hyclone),1% penicillin-streptomycin (Gibco) (D10 media for short). Cells were typically passaged every 1-2 days at a split ratio of 1:2 or 1:4 when the confluency reached at 80%.

### mRNA production by in vitro transcription and vaccine formulation

A sequence-optimized mRNA encoding B.1.351 variant (6 P) or B.1.617 variant (6P) protein was synthesized *in vitro*. The mRNA was synthesized and purified by following the manufacturer’s instructions and kept frozen at -80 °C until further use. The mRNA was encapsulated in lipid nanoparticle. The mixture was neutralized with Tris-Cl pH 7.5, sucrose was added as a cryoprotectant. The final solution was sterile filtered and stored frozen at -80 °C until further use. The particle size of mRNA-LNP was determined by DLS machine (DynaPro NanoStar, Wyatt, WDPN-06). The encapsulation and mRNA concentration were measured by using Quant-iT™ RiboGreen™ RNA Assay Kit (Thermofisher).

### Negative-stain TEM

5 µl of the sample was deposited on a glow-discharged formvar/carbon-coated copper grid (Electron Microscopy Sciences, catalog number FCF400-Cu-50), incubated for 1 min and blotted away. The grid was washed briefly with 2% (w/v) uranyl formate (Electron Microscopy Sciences, catalog number 22450) and stained for 1 min with the same uranyl formate buffer. Images were acquired using a JEOL JEM-1400 Plus microscope with an acceleration voltage of 80 kV and a bottom-mount 4k × 3k charge-coupled device camera (Advanced Microscopy Technologies, AMT).

### In vitro mRNA expression

HEK293T cells were electroporated with mRNA encoding B.1351 variant (6P) or B.1.617 variant (6P) proteins using Neon™ Transfection System 10 µL Kit following the standard protocol provided by manufacturer. After 12 h, the cells were collected and resuspended in MACS buffer (D-PBS with 2 mM EDTA and 0.5% BSA). To detect surface-protein expression, the cells were stained with 10 µg/mL ACE2–Fc chimera (Genescript, Z03484) in MACS buffer for 30 min on ice. Thereafter, cells were washed twice in MACS buffer and incubated with PE–anti-human FC antibody (Biolegend, M1310G05) in MACS buffer for 30 min on ice. Live/Dead aqua fixable stain (Invitrogen) were used to assess viability. Data acquisition was performed on BD FACSAria II Cell Sorter (BD). Analysis was performed using FlowJo software.

### Animals

*M. musculus* (mice), 6-8 weeks old females of C57BL/6Ncr, were purchased from Charles River and used for immunogenicity study. Animals were housed in individually ventilated cages in a dedicated vivarium with clean food, water, and bedding. Animals are housed with a maximum of 5 mice per cage, at regular ambient room temperature (65-75°F, or 18-23°C), 40-60% humidity, and a 14 h:10 h light cycle. All experiments utilize randomized littermate controls.

### Mice immunization and sample collection

A standard two-dose schedule given 21 days apart was adopted (Polack et al., 2020). 1 μg or 10 μg LNP-mRNA were diluted in 1X PBS and inoculated into mice intramuscularly for prime and boost. Control mice received PBS. Two weeks post-prime (day14) and two weeks post-boost (day 35), sera were collected from experimental mice and utilized for following ELISA and neutralization assay of pseudovirus. Forty days (day 40) after prime, mice were euthanized for endpoint data collection. Splenocytes were collected for T cell stimulation and cytokine analysis, and single cell profiling. Lymphocytes were separately collected from mouse blood, spleen and draining lymph nodes and applied for Bulk BCR and TCR profiling.

### Cell isolation from animals

For every mouse treated with either LNP-mRNA or PBS. Blood, spleens and draining lymph nodes were separately collected. Spleen and lymph node were homogenized gently and filtered with a 100 µm cell strainer (BD Falcon, Heidelberg, Germany). The cell suspension was centrifuged for 5 min with 400 *g* at 4 °C. Erythrocytes were lysed briefly using ACK lysis buffer (Lonza) with 1mL per spleen for 1∼2 mins before adding 10 mL PBS containing 2% FBS to restore iso-osmolarity. The single-cell suspensions were filtered through a 40 µm cell strainer (BD Falcon, Heidelberg, Germany).

### Flow Cytometry

Spleens from three mice in LNP mRNA vaccine groups and four mice in PBS group were collected five days post boost. Mononuclear single-cell suspensions from whole mouse spleens were generated using the above method. 0.5 million splenocytes were resuspended with 200µl into RPMI1640 supplemented with 10% FBS, 1% penicillin–streptomycin antibiotic, Glutamax and 2mM 2-mercaptoethonal, anti-mouse CD28 antibody (Biolegend, Clone 37.51) and seed into 96-well plate for overnight. The splenocytes were incubated for 6 hr at 37°C in vitro with BrefeldinA (Biolegend) under three conditions: no peptide, PMA/Ionomycin, and PepTivator® SARS-CoV-2 Prot_S Complete peptide pool (Miltenyi Biotec, 15 mers with 11 amino acid overlap) covering the entire SARS-CoV-2 S protein. Peptide pools were used at a final concentration of 200 ng/ml. Following stimulation, cells were washed with PBS before surface staining with LIVE/DEAD Fixable Dead Cell Stain (Invitrogen, 1:1000) and a surface stain cocktail containing the following antibodies: CD3 PE/Cy7 (Biolegend, Clone 17A2,1:200), CD8a BV421(Biolegend, Clone QA17A07, 1:200), CD4 FITC (Biolegend, Clone GK1.5,1:200) in MACS buffer (D-PBS with 2 mM EDTA and 0.5% BSA) on ice for 20 min, cells were washed with MACS buffer then fixed and permeabilized using the BD Cytofix/Cytoperm fixation/permeabilization solution kit according to the manufacturer’s instructions. Cells were washed in perm/wash solution for 5 min, and stained by intracellular staining for 30 min at 4 °C using a cocktail of the following antibodies: IFN-γ PE (Biolegend, Clone W18272D,1:500), TNF Percp-Cy5.5(Biolegend, Clone MP6-XT22, 1:500), IL2 BV510 (Biolegend, Clone JES6-5H4, 1:500), IL4 BV605 (Biolegend, Clone 11B11,1:500), IL5 APC(Biolegend, Clone TRFK5,1:500) in MACS buffer. Finally, cells were washed in MACS for twice and resuspended in MACS buffer before running on BD FACSAria II Cell Sorter (BD). Analysis was performed using FlowJo software according to the gating strategy outlined in a Supplemental Figure.

### ELISA

The 384-well ELISA plates were coated with 3 μg/ml of antigens overnight at 4 degree. The antigen panel used in the ELISA assay includes SARS-CoV-2 spike S1+S2 ECD and RBD of 2019-nCoV (SINO, ECD 40589-V08B1 and RBD 40592-V08B), Indian variant B.1.617 (SINO, ECD 40589-V08B12 and RBD 40592-V08H88), South African variant (SINO, ECD 40589-V08B07 and RBD 40592-V08H85) and. spike RBD of wild-type, South African variant and Indian variant. Plates were washed with PBS plus 0.5% Tween 20 (PBST) three times using the 50TS microplate washer (Fisher Scientific, NC0611021) and blocked with 0.5% BSA in PBST at room temperature for one hour. Plasma was serially diluted twofold or fourfold starting at a 1:2000 dilution. Samples were added to the coated plates and incubate at room temperature for one hour, followed by washes with PBST five times. Anti-mouse secondary antibody was diluted to 1:2500 in blocking buffer and incubated at room temperature for one hour. Plates were washed five times and developed with tetramethylbenzidine substrate (Biolegend, 421101). The reaction was stopped with 1 M phosphoric acid, and OD at 450 nm was determined by multimode microplate reader (PerkinElmer EnVision 2105). The binding response (OD450) were plotted against the dilution factor in log10 scale to display the dilution-dependent response. The area under curve of the dilution-dependent response (Log10 AUC) was calculated to evaluate the potency of the serum antibody binding to spike antigens.

### SARS-CoV-2 pseudovirus reporter and neutralization assays

HIV-1 based SARS-CoV-2 WT, B.1.351 variant, and B.1.617 variant pseudotyped virions were generated using respective spike sequences, and applied in neutralization assays. Plasmid expressing a C-terminally truncated SARS-CoV-2 S protein (pSARS-CoV-2Δ19) was from Dr Bieniasz’ lab. Plasmids expressing a C-terminally truncated SARS-CoV-2 B.1.351 variant S protein (B.1.351 variant-Δ19) and SARS-CoV-2 B.1.617 variant S protein (B.1.617 variant-Δ19) were generated as above. The three plasmids-based HIV-1 pseudotyped virus system were utilized to generate (HIV-1/NanoLuc2AEGFP)-SARS-CoV-2 particles, (HIV-1/NanoLuc2AEGFP)-B.1.351 variant particles, and B.1.617 variant particles. The reporter vector, pCCNanoLuc2AEGFP, and HIV-1 structural/regulatory proteins (pHIVNLGagPol) expression plasmid were gifts from Dr Bieniasz’s lab. Briefly, 293T cells were seeded in 150 mm plates, and transfected with 21 µg pHIVNLGagPol, 21 µg pCCNanoLuc2AEGFP, and 7.5 µg of a SARS-CoV-2 SΔ19 or B.1.351 variant-Δ19 or SARS-CoV-2 SA SΔ19 plasmid, utilizing 198 µl PEI. At 48 h after transfection, the 20-ml supernatant was harvested and filtered through a 0.45-µm filter, and concentrated before aliquoted and frozen in -80°C.

The pseudovirus neutralization assays were performed on 293T-hACE2 cell. One day before, 293T-hACE2 cells were plated in a 96 well plate, 0.01 x10^6^ cells per well. The following day, serial dilution serum plasma, collected from PBS or LNP-mRNA vaccine immunized mice and started from 1:100 (5-fold serial dilution using complete growth medium), 55 µL aliquots were mixed with the same volume of SARS-CoV-2 WT, B.1.351 variant, and B.1.617 variant pseudovirus. The mixture was incubated for 1 hr at 37 °C incubator, supplied with 5% CO_2_. Then 100 µL of the mixtures were added into 96-well plates with 293T-hACE2 cells. Plates were incubated at 37°C supplied with 5% CO_2_. 48 hr later, 293T-hACE2 cells were collected and the GFP+ cells were analyzed with Attune NxT Acoustic Focusing Cytometer (Thermo Fisher). The 50% inhibitory concentration (IC50) was calculated with a four-parameter logistic regression using GraphPad Prism (GraphPad Software Inc.).

### Bulk BCR and TCR sequencing

Lymphocytes from blood, draining lymph node, spleen of each mRNA-LNP vaccinated and control mice were collected as described above for mouse immunization and sample collection. mRNA of lymphocytes from three tissues were extracted using a commercial RNeasy® Plus Mini Kit (Qiagen). Following bulk BCR and TCR are prepared using SMARTer Mouse BCR IgG H/K/L Profiling Kit and SMARTer Mouse TCR a/b profiling kit separately (Takara). Based on the extracted mRNA amount of each sample, the input RNA amounts for bulk BCR libraries were as follows: lymphocytes from blood (100 ng), lymphocytes from lymph node (1000 ng), and lymphocytes from spleen (1000 ng). The input RNA amounts for bulk TCR libraries were as follows: lymphocytes from blood (100 ng), lymphocytes from lymph node (500 ng), and lymphocytes from spleen (500 ng). All procedures followed the standard protocol of the manufacture. The pooled library was sequenced using MiSeq (Illumina) with 2*300 read length.

### Bulk VDJ sequencing data analysis

Raw fastq files from bulk BCR and TCR sequencing were processed by MiXCR v2.1.5 to clonotypes. Paired-end reads were merged and aligned to reference genes for homo sapiens species using the function: mixcr align -s hs, Clones were assembled using the mixcr assemble function, then exported for specific chains (TRB, TRA, IGH, IGL, IGK) using the mixcr exportClones function. TCR-seq and BCR-seq data was subsequently analyzed using the immunarch v0.6.6 R package for clonality analyses and calculating diversity metrics such as the Chao1 estimator and Gini-Simpson index.

### Single cell profiling

Splenocytes were collected from mRNA-LNP vaccinated and control mice were collected as described above for mouse immunization and sample collection, and normalized to 1000 cells/μL. Standard volumes of cell suspension were loaded to achieve targeted cell recovery to 10000 cells. The samples were subjected to 14 cycles of cDNA amplification. Following this, gene expression (GEX), TCR-enriched and BCR-enriched libraries were prepared according to the manufacturer’s protocol (10x Genomics). All libraries were sequenced using a NovaSeq 6000 (Illumina) with 2*150 read length.

### Single cell transcriptomics data analysis for immune repertoire profiling

Both standard established pipelines and custom scripts were used for processing and analyzing single cell GEX data. Illumina sequencing data was processed using the Cellranger v5.0.1 (10x Genomics) pipeline and aligned to the mm10 reference. Cellranger outputs were then processed and analyzed using standard Seurat v. 4.0.2 workflow, including log normalization with scale factor 10,000, scaling and centering, principal components analyses, nearest-neighbor graph construction, clustering with the Louvain algorithm, uniform manifold approximation and projection (UMAP), differential gene expression, and generation of various visualizations. The following parameters were used: for the FindNeighbors function, dims = 1:10; for FindClusters, resolution = 0.6; for RunUMAP, dims = 1:10; for FindAllMarkers, only.pos = TRUE, min.pct = 0.25, logfc.threshold = 0.25.

Assignment of immune cell type identity to clusters was performed manually based on expression of cell type specific markers. Custom scripts were used for cell proportion calculations and condition-specific analyses and statistics (e.g. Wilcoxon rank sum test). While cluster 5 cells were annotated as “CD8 T / NKT cell” as it was a mixed population, these cells were merged with the “CD8 T cell” annotation for proportion calculations after cells with greater than 1 expression for any of the following markers were removed: *Klrb1, Klra6, Klra1, Zbtb16*. Differential gene expression between conditions for various cell types were performed by the FindMarkers function with the parameters logfc.threshold = 0.01 and min.pct = 0.1. For T-cell specific analyses, cells associated with the following terms were taken as a subset and used for standard Seurat pipeline analyses as described above: “CD4 T cell”, “CD8 T / NKT cell”, “CD8 T cell”, “T cell-like.” For B-cell specific analyses, cells associated with the following terms were taken as a subset: “B cell”, “B cell-like”, “Progenitor B cell”, “Plasma cell.”

For functional annotation, differentially upregulated and downregulated genes with cutoff of adjusted p-value 0.05 were used for DAVID analysis. Genes associated with gene ontology terms “regulation of immune effector process” (GO:0002697), “immune response” (GO:0006955), “regulation of T cell activation” (GO:0050863), and “regulation of B cell activation” (GO:0050864) were used for generating annotation-associated heatmaps. Custom R scripts were used for generating various other figures.

### Single cell VDJ sequencing data analysis

Illumina sequencing data was processed using the Cellranger v5.0.1 (10x Genomics) pipeline. The filtered_contig_annotations output file was used as an input to immunarch v0.6.6 R package for calculating diversity metrics such as the Chao1 estimator and Gini-Simpson index. The clonotypes output file was used for analysis with custom scripts for clonality analyses and CDR3 distribution ring plots.

### Statistical analysis

The statistical methods are described in figure legends and/or supplementary Excel tables. The statistical significance was labeled as follows: n.s., not significant; * p < 0.05; ** p < 0.01; *** p < 0.001; **** p < 0.0001. Prism (GraphPad Software) and RStudio were used for these analyses. Additional information can be found in the Nature Research Reporting Summary.

### Replication, randomization, blinding and reagent validations

Replicate experiments have been performed for all key data shown in this study.

Biological or technical replicate samples were randomized where appropriate. In animal experiments, mice were randomized by littermates.

Experiments were not blinded.

NGS data processing were blinded using metadata. Subsequent analyses were not blinded.

Commercial antibodies were validated by the vendors, and re-validated in house as appropriate. Custom antibodies were validated by specific antibody - antigen interaction assays, such as ELISA. Isotype controls were used for antibody validations.

Cell lines were authenticated by original vendors, and re-validated in lab as appropriate.

All cell lines tested negative for mycoplasma.

## Data, resources and code availability

All data generated or analyzed during this study are included in this article and its supplementary information files. Specifically, source data and statistics for non-high-throughput experiments are provided in a supplementary table excel file. High-throughput experiment data are provided as processed quantifications in Supplemental Datasets. Genomic sequencing raw data are deposited to Gene Expression Omnibus (GEO) with a pending accession code. Codes that support the findings of this research are being deposited to a public repository such as GitHub. Additional information related to this study are available from the corresponding authors upon reasonable request.

---

**Figure S 1.**
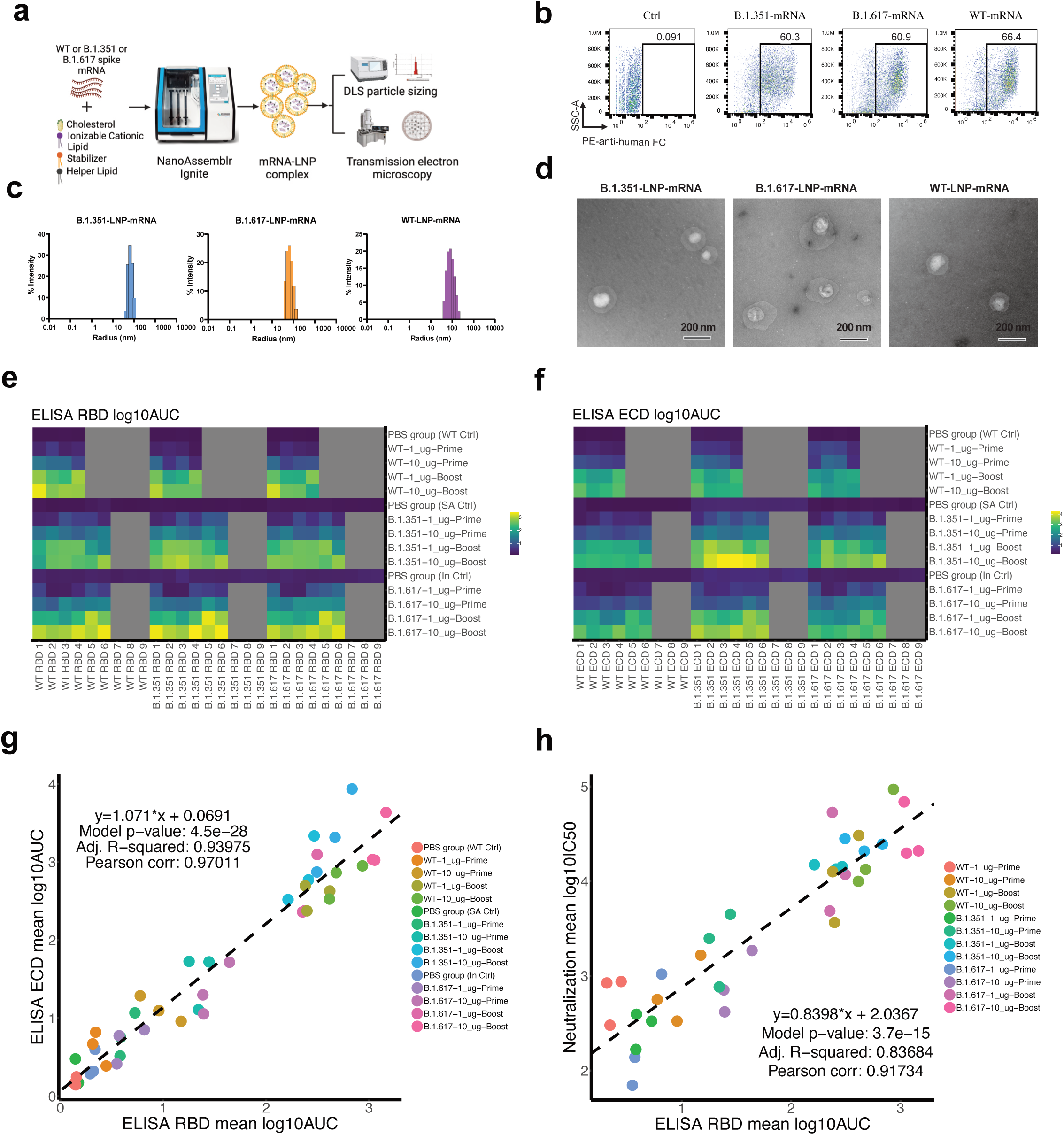
Workflow of production and physical characterization of the spike mRNA and lipid nanoparticles used for the LNP-mRNA vaccine candidates; and additional data on antibody responses of all three groups of LNP-mRNA vaccinated animals. **a**, Graphical representation of B.1.351-LNP-mRNA complex and B.1.617-LNP-mRNA complex formation. The spike mRNAs of B.1.351 and B.1.617 were encapsulated by LNP via NanoAssemblr Ignite. The size and encapsulation rate of the mRNA-LNP complex were measured by dynamic light scatter (DLS) and Ribogreen assay, respectively. **b**, After electroporated into 293FT cells, *In vitro* expression of B.1.351-spike or B.1.617-spike mRNA were detected by flowcytometry using the human ACE2-Fc fusion protein and PE-anti-Fc antibody. **c-d**, DLS (c) and TEM (d) of size and monodispersity characterization of LNP-mRNAs. **e**, Heatmap of ELISA titers of animals vaccinated by all three LNP-mRNAs, against the RBDs from three different spikes of SARS-CoV-2. **f**, Heatmap of ELISA titers of animals vaccinated by all three LNP-mRNAs, against the ECDs from three different spikes of SARS-CoV-2. **g**, Correlation X-Y scatterplots of ELISA and neutralization titers between ELISA RBD log10AUC vs ELISA ECD log10AUC for all vaccine groups **h**, Correlation X-Y scatterplots of ELISA and neutralization titers between ELISA RBD log10AUC vs neutralization log10IC50 for all vaccine groups. **Related to: Figure(s) 1, 2**

**Figure S 2.**
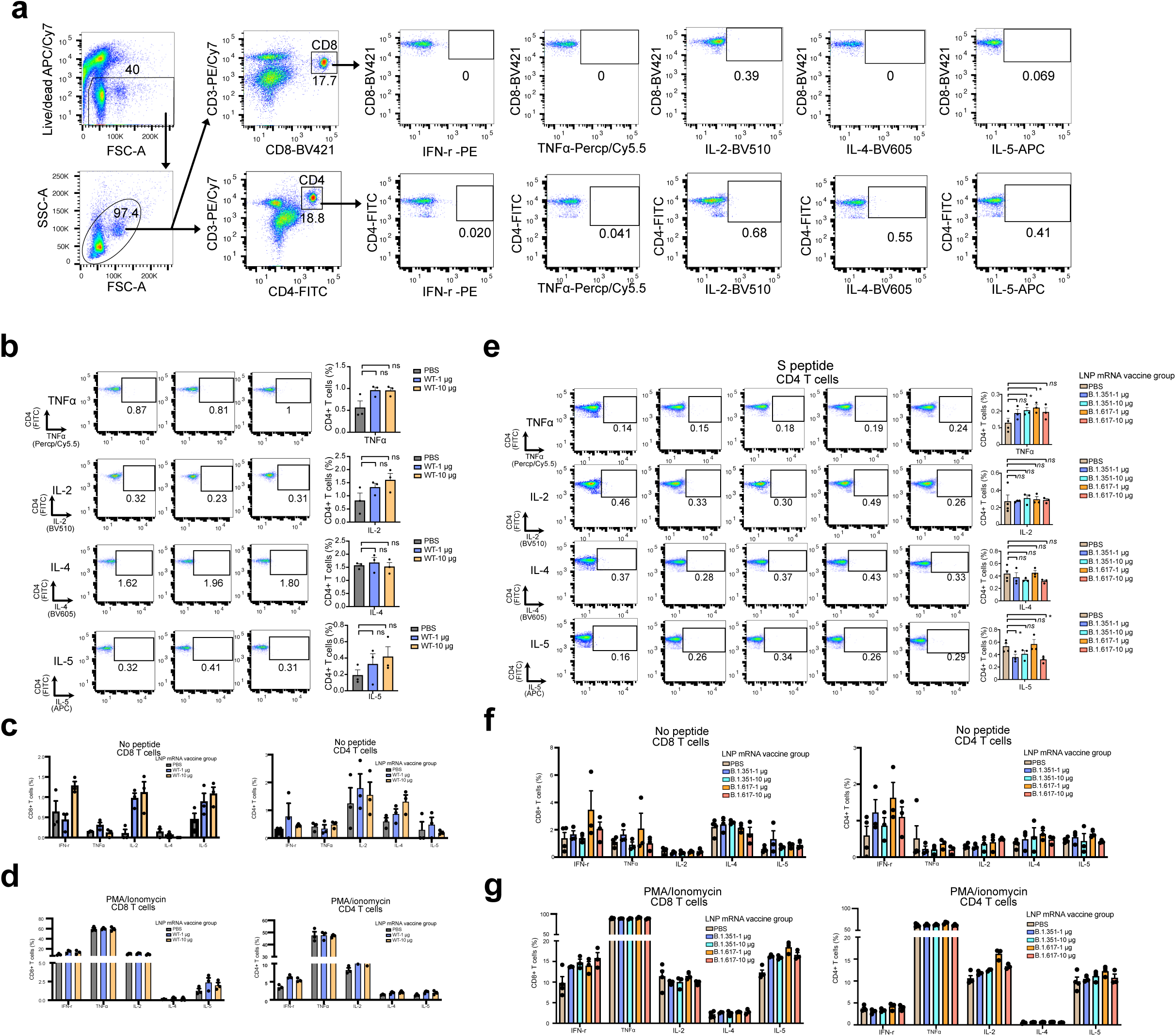
Additional flow cytometry analysis of WT-LNP-mRNA, B.1.351-LNP-mRNA and B.1.617-LNP-mRNA induced S protein-specific T cell response. **a**, Flow cytometry panel and gating strategy to quantify SARS-CoV-2 S-specific T cells in B.1.351-mRNA-LNP and B.1.617-mRNA-LNP vaccinated group and PBS group. **b**, Percentage of CD4 T cells expressing TNFα, IL-2, IL4, and IL5 in response to S peptide pools of WT-mRNA-LNP vaccines treated mice and PBS treated control mice. **c-d**, Percentage of CD8 T cells (left) and CD4 cells (right) expressing IFN-γ, TNFα, IL-2, IL-4, and IL-5 of splenocytes from WT-LNP-mRNA vaccinated mice and PBS treated control mice without peptide stimulation (c), or in response to PMA/ionomycin stimulation (d). **e**, Percentage of CD4 T cells expressing TNFα, IL-2, IL4, and IL5 in response to S peptide pools of B.1.351-mRNA-LNP and B.1.617-mRNA-LNP vaccinated group and PBS group. **f-g**, Percentage of CD8 T cells (left) and CD4 cells (right) expressing IFN-γ, TNFα, IL-2, IL-4, and IL-5 of splenocytes from B.1.351-mRNA-LNP and B.1.617-mRNA-LNP vaccinated mice and PBS treated control mice without peptide stimulation (f), or in response to PMA/ionomycin stimulation (g). **Notes:** In this figure: Each dot represents data from one mouse. Statistical significance labels: n.s., not significant; * p < 0.05; ** p < 0.01; *** p < 0.001; **** p < 0.0001 **Related to: Figure(s) 1, 3**

**Figure S 3.**
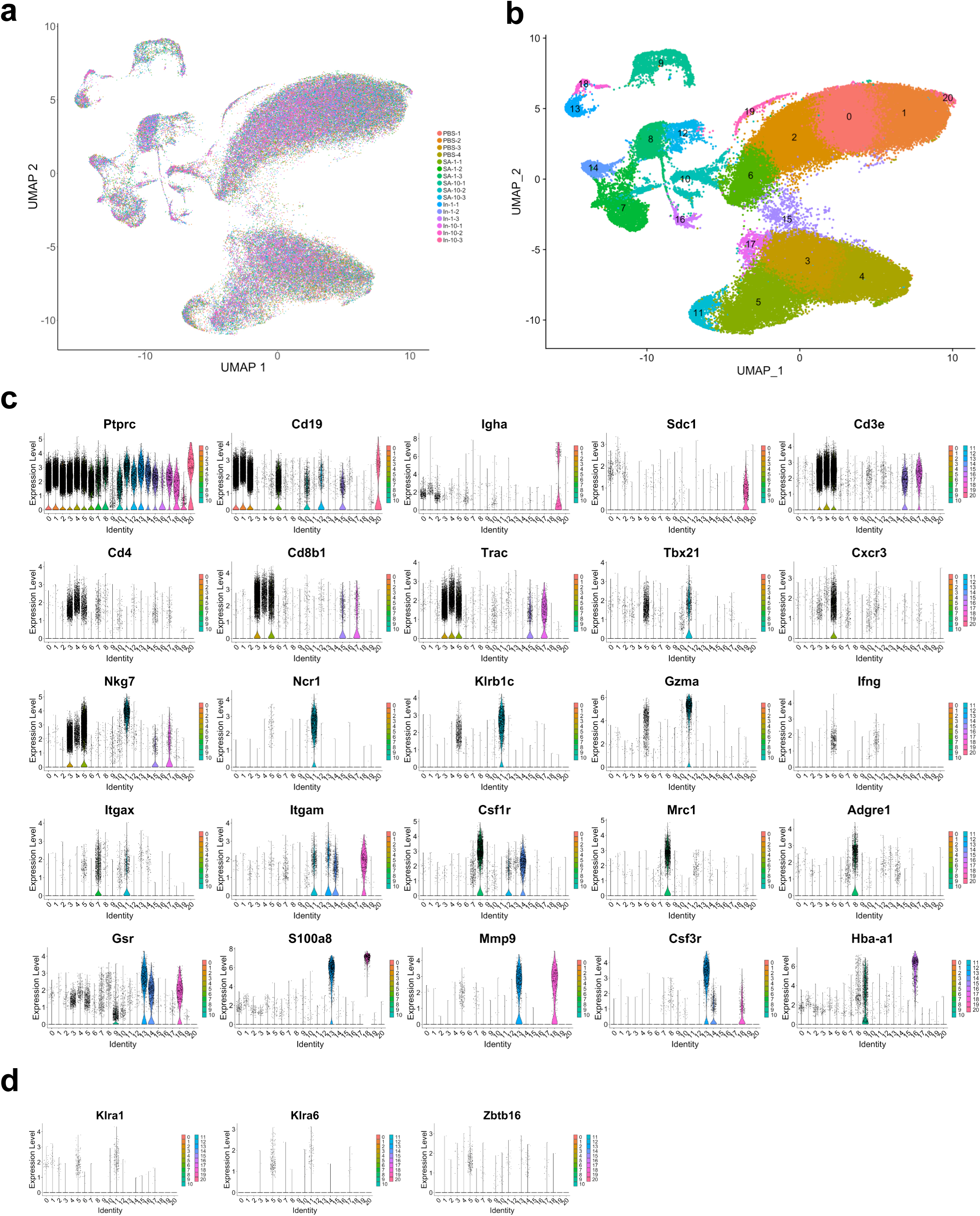
Additional analysis of single cell transcriptomics and immune repertoire profiling of B.1.351-LNP-mRNA and B.1.617-LNP-mRNA vaccinated animals. **a,** UMAP visualization of all 90,152 cells pooled across samples and conditions. Cells colored by sample. **b,** UMAP visualization of all 90,152 cells pooled across samples and conditions. Cells colored and labelled by cluster. **c,** Violin plots of expression of representative cell type specific markers used for assignment of cell identity. **d,** Violin plots of expression of representative cell type specific markers (Klra1, Klra6, and Zbtb16) used for denoting the existence of NKT cells within a CD8 T cell dominant cluster (Cluster 5 of overall UMAP). **Related to: Figure(s) 4**

**Figure S 4.**
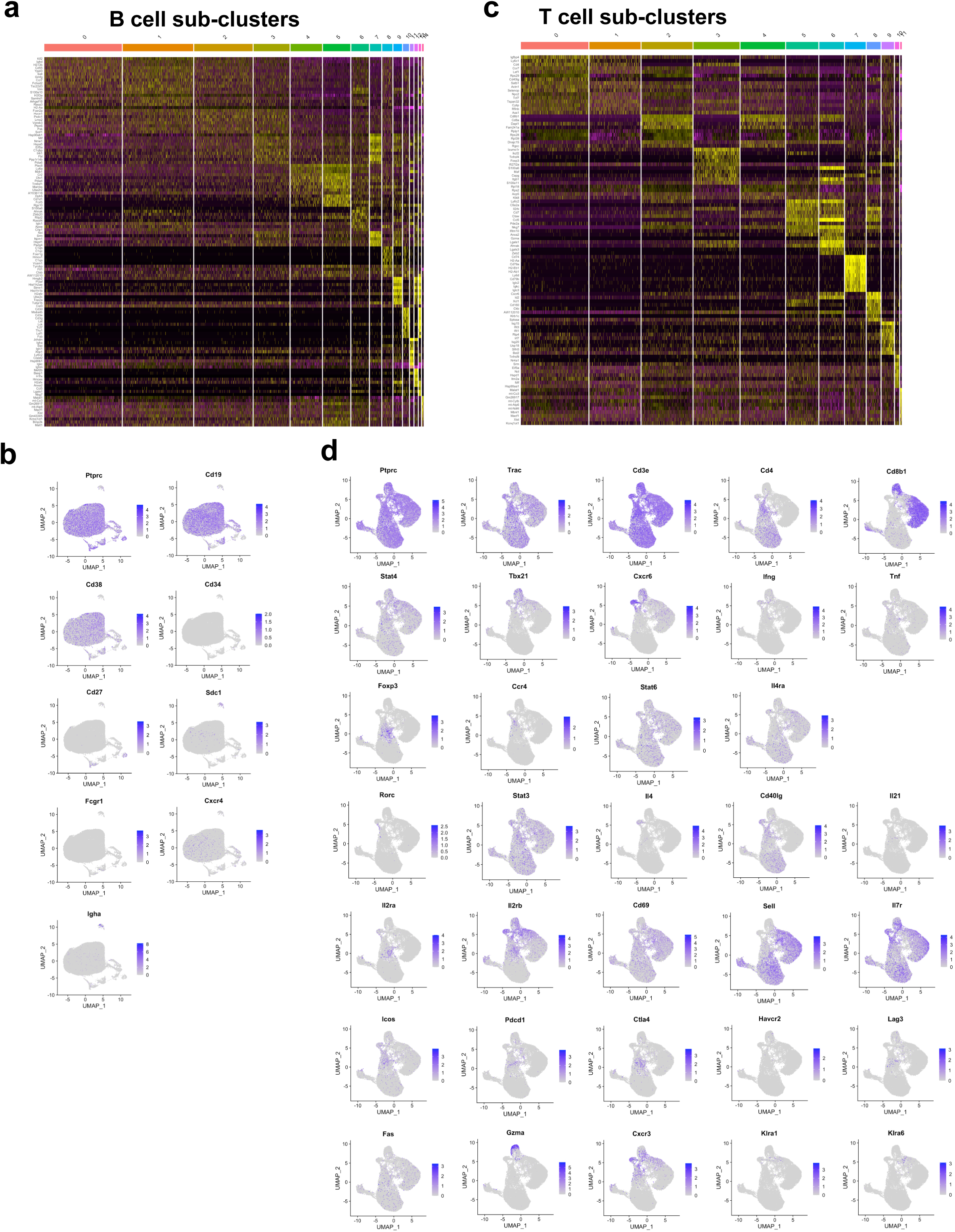
Additional analysis B cell and T cell subpopulations in scRNA-seq profiling of variant-specific LNP-mRNA vaccinated animals. **a,** Heatmap of differentially expressed markers for all sub-clusters (n = 15) from B cell-associated subpopulation UMAP and clustering. **b,** UMAP visualizations highlighting expression levels of individual representative marker genes from B cell-associated subpopulation analyses. **c,** Heatmap of differentially expressed markers for all sub-clusters (n = 2) from T cell-associated subpopulation UMAP and clustering. **d,** UMAP visualizations highlighting expression levels of individual genes from T cell-associated subpopulation analyses. Representative marker categorizations include: Th1, Th2, Treg, Th17, Tfh, activation, memory, co-stimulation, exhaustion, effector function, and NKT. **Related to: Figure(s) 5**

**Figure S 5.**
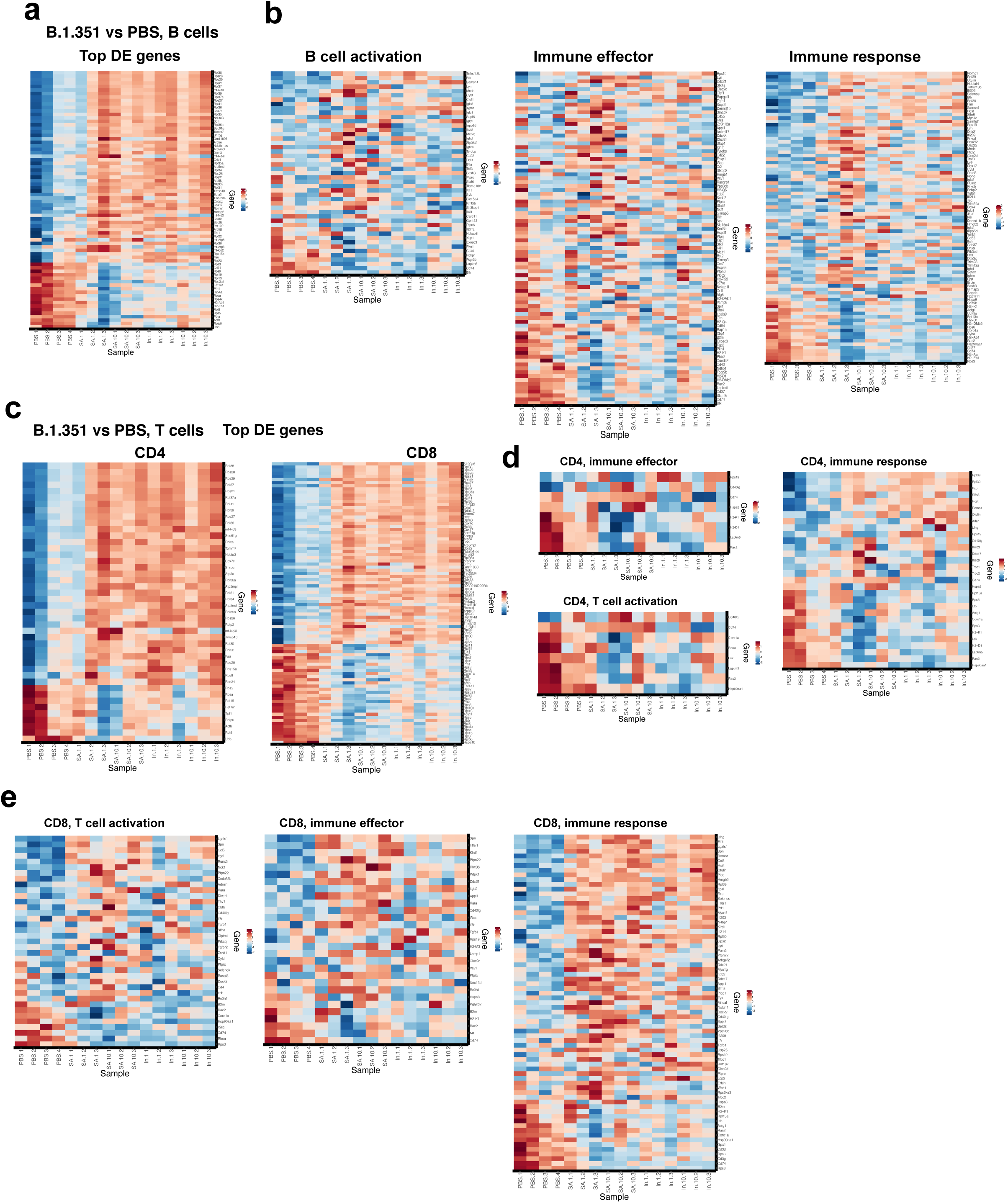
Additional analysis of differential gene expression in the B and T cell subpopulations of variant-specific LNP-mRNA vaccinated animals. **a,** Heatmap of top differentially expressed genes between B.1.351-LNP-mRNA sample B cells vs PBS sample B cells, differential expression profiles shown for all samples (n = 16). **b,** Heatmap of B cell differential expression profiles with intersection of immune genes. Markers taken from the intersection of genes associated with the “regulation of B cell activation”, “immune effector” or “immune response” Gene Ontology annotations and differentially expressed genes between B.1.351-LNP-mRNA sample B cells vs PBS sample B cells and B.1.617-LNP-mRNA sample B cells vs PBS sample B cells (intersect between the GO genes and the union set of DE genes). **c,** Heatmaps of top differentially expressed genes between B.1.351-LNP-mRNA sample CD4 T cells vs PBS sample CD4 T cells and B.1.351-LNP-mRNA sample CD8 T cells vs PBS sample CD9 T cells, expression profiles shown for all samples (n = 16) for each heatmap. **d-e,** Heatmaps of CD4 (**d**) and CD8 (**e**) T cell differential expression profiles with intersection of immune genes. Markers taken from the intersection of genes associated with the “regulation of T cell activation”, “immune effector” or “immune response” Gene Ontology annotations and differentially expressed genes between B.1.351-LNP-mRNA sample T cells vs PBS sample T cells and B.1.617-LNP-mRNA sample T cells vs PBS sample T cells (intersect between the GO genes and the union set of DE genes), for respective CD4 and CD8 T cell comparisons. **Related to: Figure(s) 5**

**Figure S 6.**
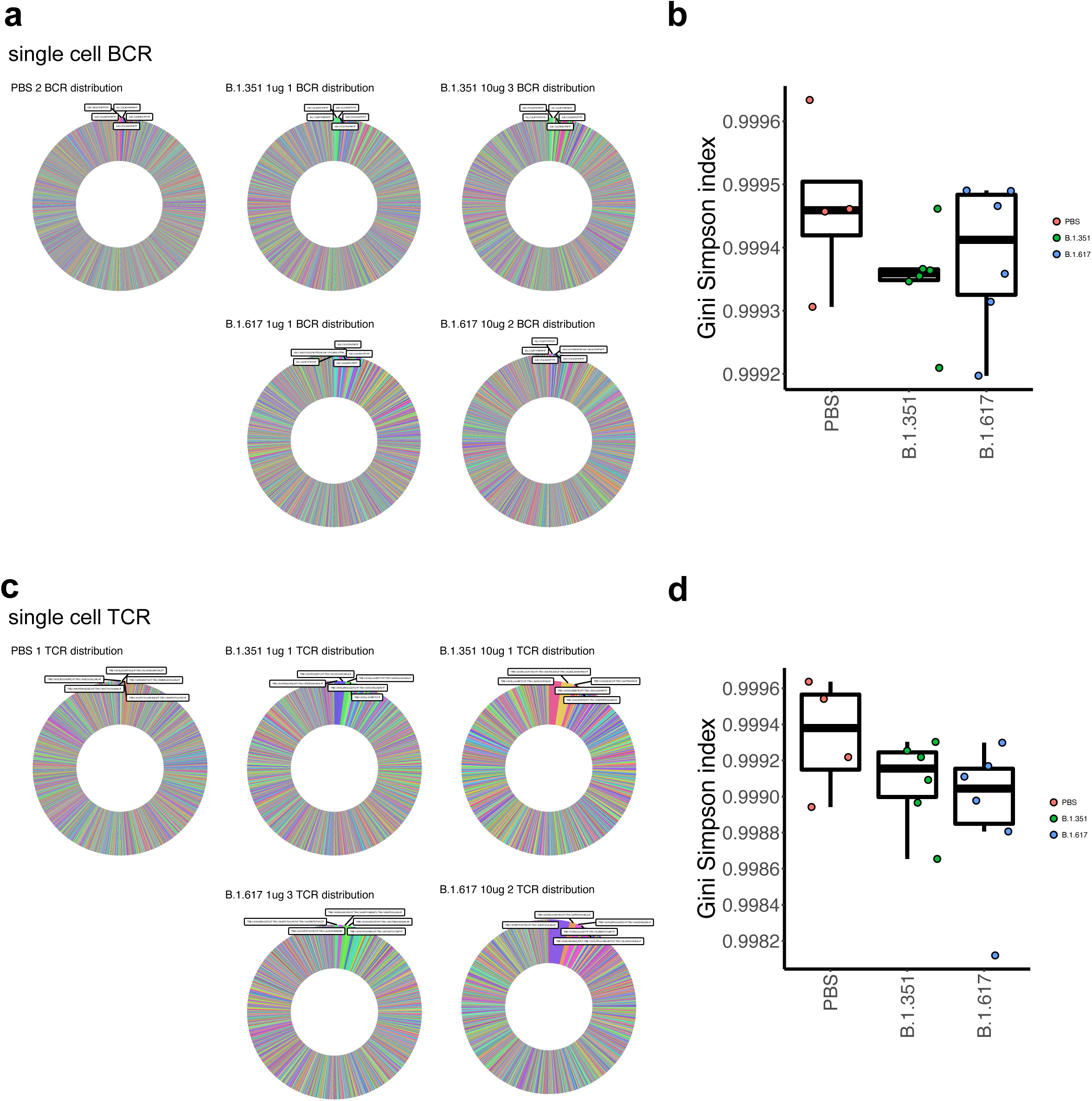
Additional analysis of single cell VDJ BCR-seq and TCR-seq. **a,** Ring plots of BCR distribution for single cell BCR-seq dataset. Top BCR sequences were labeled in the plot. **b,** Boxplot of Gini-Simpson indices for each condition (PBS, n = 4; B.1.351-LNP-mRNA, n = 6; B.1.617-LNP-mRNA, n = 6) for repertoires in the single cell BCR-seq dataset. **c,** Ring plots of TCR distribution for single cell TCR-seq dataset. Top TCR sequences were labeled in the plot. **d,** Boxplot of Gini-Simpson indices for each condition (PBS, n = 4; B.1.351-LNP-mRNA, n = 6; B.1.617-LNP-mRNA, n = 6) for repertoires in the single cell TCR-seq dataset. **Related to: Figure(s) 6**

**Figure S 7.**
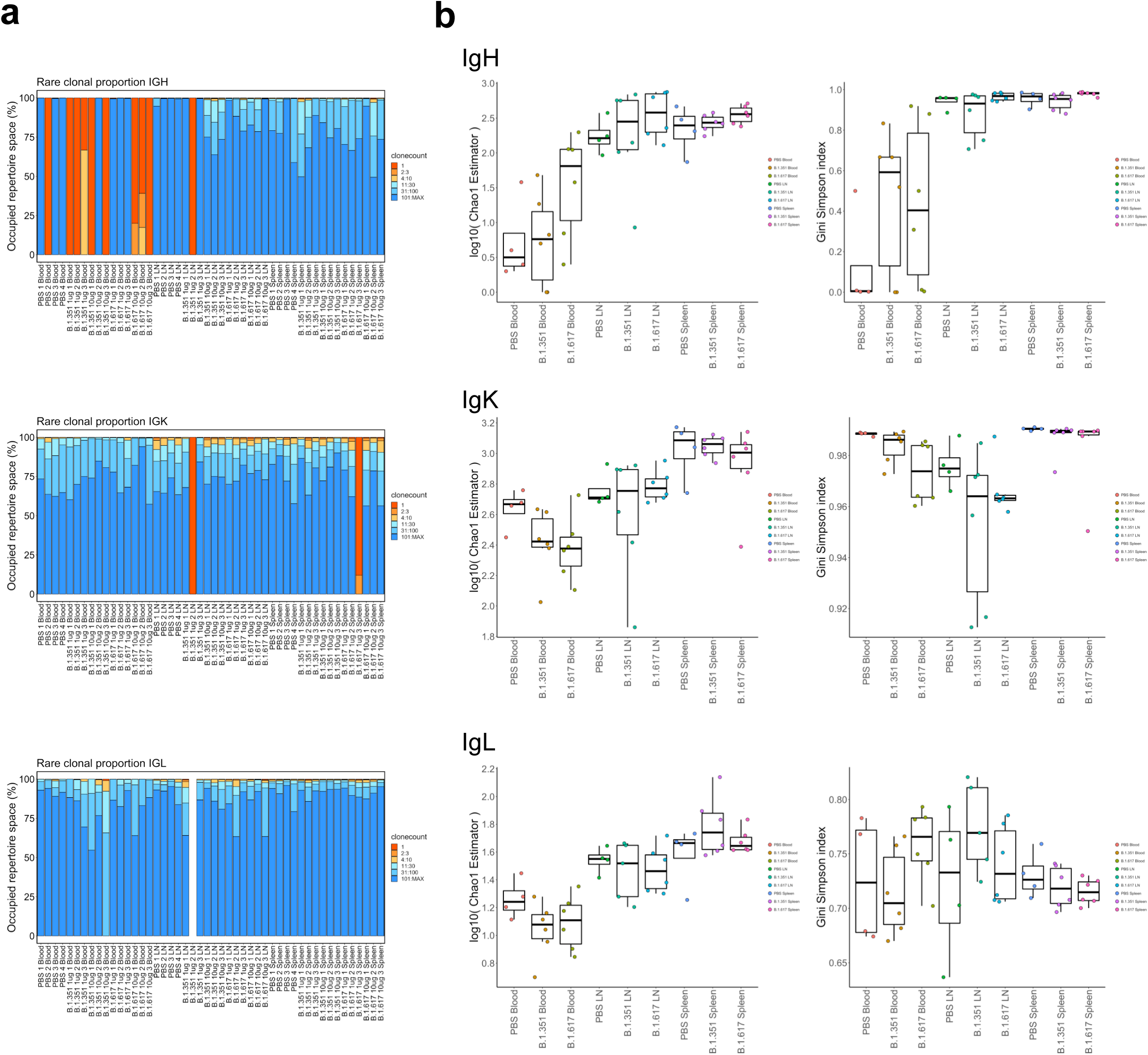
Additional analysis of bulk VDJ BCR-seq and TCR-seq. **a,** Clonotypes bar plot depicting rare clone proportions of IgH, IgK or IgL chains for all samples (n = 48) in the bulk BCR-seq dataset. **b,** Boxplots of Chao1 (left) and Gini-Simpson (right) indices of clonality of IgH, IgK or IgL chains across vaccination and tissue of origin groups. Noted the low dose and high dose groups of the same vaccine were grouped together. **Related to: Figure(s) 6**

